# Different complex regulatory phenotypes underlie hybrid male sterility in divergent rodent crosses

**DOI:** 10.1101/2023.10.30.564782

**Authors:** Kelsie E. Hunnicutt, Colin Callahan, Sara Keeble, Emily C. Moore, Jeffrey M. Good, Erica L. Larson

## Abstract

Hybrid incompatibilities are a critical component of species barriers and may arise due to negative interactions between divergent regulatory elements in parental species. We used a comparative approach to identify common themes in the regulatory phenotypes associated with hybrid male sterility in two divergent rodent crosses, dwarf hamsters and house mice. We investigated three potential characteristic gene expression phenotypes in hybrids including the propensity of transgressive differentially expressed genes towards over or underexpression, the influence of developmental stage on patterns of misexpression, and the role of the sex chromosomes on misexpression phenotypes. In contrast to near pervasive overexpression in hybrid house mice, we found that misexpression in hybrid dwarf hamsters was dependent on developmental stage. In both house mouse and dwarf hamster hybrids, however, misexpression increased with the progression of spermatogenesis, although to varying extents and with potentially different consequences. In both systems, we detected sex-chromosome specific overexpression in stages of spermatogenesis where inactivated X chromosome expression was expected, but the hybrid overexpression phenotypes were fundamentally different. Importantly, misexpression phenotypes support the presence of multiple developmental blocks to spermatogenesis in dwarf hamster hybrids, including a potential role of meiotic stalling or breakdown early in spermatogenesis. Collectively, we demonstrate that while there are some similarities in hybrid expression phenotypes of house mice and dwarf hamsters, there are also clear differences that point towards unique mechanisms underlying hybrid male sterility. Our results highlight the potential of comparative approaches in helping to understand the causes and consequences of disrupted gene expression in speciation.

## INTRODUCTION

The evolution of postzygotic reproductive barriers, such as hybrid inviability and sterility, is an important part of the speciation process, and identifying the genetic architecture of hybrid incompatibilities has been a common goal uniting speciation research (Coughlan and Matute 2020). While identifying the genetic basis of hybrid dysfunction remains difficult in many systems, downstream regulatory phenotypes can provide insight into the underlying mechanisms of speciation (Mack and Nachman 2017). An outstanding question surrounding the role of disrupted gene regulation and speciation is whether the combination of two divergent genomes in hybrids results in gene expression perturbations that are consistent or repeatable across species. At the broadest level, it is unclear whether gene expression in hybrids tends to be intermediate or transgressive (outside the range of parental gene expression), whether transgressive gene expression is biased towards over or underexpression (Ortíz-Barrientos *et al*. 2007), and whether transgressive misexpression tends to be modulated by *cis* or *trans* regulatory elements (Wittkopp *et al*. 2004; McManus *et al*. 2010; Oka *et al*. 2014; Mack *et al*. 2016; Mugal *et al*. 2020; Kopania *et al*. 2022a). By investigating trends in the magnitude and direction of transgressive expression across different hybrid systems, we can begin to understand the evolutionary forces shaping regulatory-based hybrid incompatibilities. For example, if transgressive misexpression in hybrids tends towards overexpression, this may mean that genes with disrupted regulation in hybrids tend to be genes that are normally repressed in parental lineages (Meiklejohn *et al*. 2014; Barreto *et al*. 2015; Larson *et al*. 2017). Alternatively, transgressive misexpression in hybrids may tend towards underexpression if regulatory divergence between parental lineages results in impaired transcription factor binding with promoter or enhancer elements (Oka *et al*. 2014; Guerrero *et al*. 2016) or if divergence stimulates epigenetic silencing (Paun *et al*. 2007; Shivaprasad *et al*. 2012; Lafon-Placette and Köhler 2015; Brekke *et al*. 2016; Zhu *et al*. 2017). At a finer scale, transgressive misexpression patterns may depend on developmental stage: for example, if there is greater pleiotropy earlier in development (Ortíz-Barrientos *et al*. 2007; Cutter and Bundus 2020). In particular, we might expect sterile hybrids to have more transgressive misexpression during later stages of gametogenesis when genes are evolving rapidly and are potentially under less regulatory constraint (Kopania *et al*. 2022a; Murat *et al*. 2023). Finally, the role of sex chromosome regulation in inviable or sterile hybrids encompasses both larger questions. Sex chromosomes may be prone to asymmetry in their expression divergence (Oka and Shiroishi 2014; Civetta 2016) and be regulated differently across stages of development (Presgraves 2008; Larson *et al*. 2018), particularly in reproductive tissues, and thus may play a central role in hybrid dysregulation relative to autosomes.

Disruption of sex chromosome regulation is thought to be a potentially widespread regulatory phenotype in sterile hybrids (Lifschytz and Lindsley 1972; Larson *et al*. 2018), in part because X chromosome repression may be crucial to normal spermatogenesis in diverse taxa (McKee and Handel 1993; Landeen *et al*. 2016; Taxiarchi *et al*. 2019; Rappaport *et al*. 2021; Viera *et al*. 2021; Murat *et al*. 2023). Furthermore, misregulation of the X chromosome is associated with hybrid sterility in several species pairs (Davis *et al*. 2015; Sánchez-Ramírez *et al*. 2021; Bredemeyer *et al*. 2021), although it has been best studied in house mice. In fertile male mice, the X chromosome is silenced just prior to the Diplotene stage of meiosis through meiotic sex chromosome inactivation (MSCI; McKee and Handel 1993; Handel 2004) and is again repressed in postmeiotic sperm development (*i.e.,* postmeiotic sex chromosome repression or PSCR; Namekawa *et al*. 2006). In contrast, the X chromosome is not properly inactivated and is overexpressed in sterile hybrid mice (Good *et al*. 2010; Bhattacharyya *et al*. 2013; Campbell *et al*. 2013; Turner *et al*. 2014; Larson *et al*. 2017, 2022). Disrupted MSCI in house mice is associated with divergence at *Prdm9*, a gene that is a major contributor to hybrid male sterility (Mihola *et al*. 2009; Davies *et al*. 2016). However, misexpression of the X chromosome in sterile hybrids could result from mechanisms other than *Prdm9*-associated disrupted MSCI, such as mispairing of the sex chromosomes due to divergence in their region of homology known as the pseudoautosomal region (PAR; Burgoyne 1982; Ellis and Goodfellow 1989; Raudsepp and Chowdhary 2015). In sum, the ubiquity of X chromosome repression and the growing body of evidence linking disrupted MSCI to hybrid sterility in mammals suggest that disrupted sex chromosome regulation may be a common regulatory phenotype in sterile hybrid males.

Here, we characterized disruption of gene expression associated with hybrid male sterility in two rodent crosses, dwarf hamsters and house mice, which span ∼35 million years of divergence (Swanson *et al*. 2019). The regulatory phenotypes of hybrid male sterility have been thoroughly studied in house mice (Good *et al*. 2010; Bhattacharyya *et al*. 2013; Campbell *et al*. 2013; Turner *et al*. 2014; Larson *et al*. 2017, 2022; Hunnicutt *et al*. 2022). We contrast these with an analogous cross between two sister species of dwarf hamster, Campbell’s dwarf hamster (*Phodopus campbelli*) and the Siberian dwarf hamster (*P. sungorus*), and their sterile F1 hybrid male offspring. These species diverged only ∼0.8-1.0 million years ago (Neumann *et al*. 2006), and they are not thought to interbreed in the wild due to geographic separation (Ishishita *et al*. 2015). Crosses between female *P. sungorus* and male *P. campbelli* produce sterile hybrid males that, similar to mice, have a range of sterility phenotypes, suggesting multiple developmental blocks to spermatogenesis (Ishishita *et al*. 2015; Bikchurina *et al*. 2018). Hybrids from the reciprocal cross are usually inviable due to abnormal growth *in utero* (Brekke and Good 2014), and the species origin of the X chromosome is the primary genetic factor controlling hybrid inviability (Brekke *et al*. 2021). Additionally, sex chromosome asynapsis during spermatogenesis is common in hybrid dwarf hamsters, providing further reason to think that X chromosome-specific misregulation may be observed in sterile hybrid dwarf hamsters (Ishishita *et al*. 2015; Bikchurina *et al*. 2018). Both the abnormal spermatogenic phenotypes observed in dwarf hamster hybrids and the potential regulatory interactions that may result from the involvement of the X chromosome in multiple reproductive barriers make dwarf hamsters an important comparison to mice for investigating what regulatory phenotypes may be repeatedly associated with the evolution of postzygotic reproductive isolation.

Expression phenotypes associated with hybrid sterility have historically been difficult to assess because of the cellular diversity of reproductive tissues (*e.g.,* testes; Ramm and Schärer 2014) and because hybrids may differ from parents in both tissue composition and developmental timing (reviewed in Montgomery and Mank 2016; Hunnicutt *et al*. 2022). To overcome these difficulties, we used Fluorescence Activated Cell Sorting (FACS) to isolate and sequence cell populations across the developmental timeline of spermatogenesis for each species pair and their F1 hybrids, including stages that span the different sex chromosome regulatory states. Our developmental timeline spans stages where we expect fertile parents to have a transcriptionally active X chromosome (spermatogonia and leptotene/zygotene spermatocytes) and an inactive X chromosome (diplotene spermatocytes and round spermatids). We used both datasets to address three main questions about the transgressive gene expression phenotypes observed in sterile hybrids: (1) within transgressive differentially expressed genes, does misexpression tend towards up- or downregulation in hybrids compared to parents? (2) are there similar patterns of disrupted transgressive expression across stages of development? and (3) are there clear differences between autosomes and sex chromosomes in expression phenotypes? And if so, is sex chromosome-specific transgressive misexpression consistent with either disrupted MSCI and/or disrupted PAR regulation? Collectively, we demonstrate the power of cell type-specific approaches for untangling the expression phenotypes associated with the evolution of hybrid male sterility and for identifying common themes in the mechanistic basis of hybrid incompatibilities across divergent taxa.

## MATERIALS AND METHODS

### Hamster crosses and male reproductive phenotypes

We used wild-derived colonies of two sister species of dwarf hamster, *P. sungorus* and *P. campbelli*, established by Kathy Wynne-Edwards (Scribner and Wynne-Edwards 1994) and housed at the University of Montana. Both species were maintained as closed colonies with a breeding scheme to minimize inbreeding. Nonetheless, inbreeding levels of these closed colonies are still high as indicated by very low nucleotide diversity (Brekke *et al*. 2018). We used males from both parent species and male F1 hybrid offspring from crosses of female *P. campbelli* with male *P. sungorus*. We weaned males in same-sex sibling groups between 17 - 21 dpp and housed them individually at 45 dpp. We euthanized reproductively mature males using carbon dioxide followed by cervical dislocation between 59 - 200 dpp (Table S1 in File S1). All animal use was approved by the University of Montana (IACUC protocols 050-16JGDBS & 035-19JGDBS).

We measured several fertility metrics for parent species and hybrid males including paired testes weight, paired seminal vesicle weight, normalized sperm counts, and sperm motility (Good *et al*. 2008). Paired testes weight and paired seminal vesicle weight were correlated with body weight (paired testes weight Pearson’s *r*(29) = 0.47, p = 0.007; paired seminal vesicle weight Pearson’s *r*(23) = 0.56, p = 0.003), so we standardized both metrics relative to body weight. We calculated sperm count by isolating sperm from caudal epididymides diced in 1 ml of Dulbecco’s PBS (Sigma) and incubated at 37°C for 10 minutes. We quantified sperm motility (proportion of motile sperm in a 5 µl suspension) and sperm count (number of sperm with head and tail in a heat shocked 5 µl suspension) across a fixed area on a Makler counting chamber. We performed statistical comparisons of fertility phenotypes in R v.4.3.1, and we used the FSA package v.0.9.4 for the Kruskal-Wallis and Dunn’s tests (Ogle and Ogle 2017).

### Isolation of enriched cell populations from hamster testes

To investigate the regulatory dynamics of the sex chromosomes during spermatogenesis, we isolated four spermatogenic cell populations from whole testes using FACS. These cell populations span a developmental timeline of spermatogenesis from mitosis (spermatogonia), meiosis prior to X inactivation (leptotene/zygotene spermatocytes), meiosis, after X inactivation (diplotene spermatocytes), and post-meiosis (round spermatids). Briefly, we disassociated a single testis per male following a published protocol originally developed for house mice (Getun *et al*. 2011) with modifications (github.com/goodest-goodlab/good-protocols/tree/main/protocols/FACS, last accessed June 16, 2021). We doubled the volumes of all reagents to account for the increased mass of testes in dwarf hamsters relative to house mice. We isolated cell populations based on size, granularity, and fluorescence on a FACSAria IIu cell sorter (BD Biosciences) at the University of Montana Center for Environmental Health Sciences Fluorescence Cytometry Core. For each sorted cell population, we extracted RNA using RNeasy kits (Qiagen) following protocols for Purification of Total RNA from Animal Cells. We quantified sample RNA quantity and quality (requiring an RNA integrity number > 7) on a Tapestation 2200 (Agilent) at the University of Montana genomics core. RNA libraries were prepared by Novogene and sequenced on Illumina NovaSeq 6000s (paired end, 150 bp). Six samples (distributed across different species and cell types) had low RNA concentrations, and for these samples we used Novogene’s low input RNA library preparation (Table S2 in File S1). MDS plots indicated no severe library batch effects between samples from different library preparations (Figure S1 in File S1), so we included all samples from both libraries in subsequent analyses.

### Read processing and mapping

We sequenced RNA from each cell population for three to five individuals of each parent species and F1 hybrids generating an average of ∼27.5 million read pairs per individual (Table S2 in File S1). We trimmed reads using Trimmomatic v.0.39 (Bolger *et al*. 2014) to remove low quality bases from the first and last 5 bp of each read and bases with an average Phred score of less than 15 across a 4 bp sliding window and only retained reads of at least 36 bp. We next used an approach (based on the modtools pipeline) which maps reads from each sample to pseudogenomes for both parent species (described below) to obtain a merged output alignment file in order to alleviate reference bias associated with mapping hybrids to only a single reference genome (Holt *et al*. 2013; Huang *et al*. 2014). For this approach, we mapped reads for each individual to both a *P. sungorus* pseudogenome and a *P. campbelli* pseudogenome with Hisat v.2.2.0 (Kim *et al*. 2019) with default settings and retaining at most 100 distinct, primary alignments, although multi-mapped reads were removed downstream (described below). We generated the *P. sungorus* pseudogenome by mapping RNASeq reads from a male *P. sungorus* individual (30.6 million total read pairs; NCBI SRA: SRR17223284; Moore *et al*. 2022) to the *P. sungorus* reference genome (GCA_023856395.1) with bwa-mem v.2.2.1 (Vasimuddin *et al*. 2019), and the *P. campbelli* pseudogenome by mapping female *P. campbelli* whole genome sequencing reads (average coverage: 33x; NCBI SRA: SRR17223279; Moore *et al*. 2022) to the *P. sungorus* reference genome. Because our *P. sungorus* pseudogenome was based on a male hamster and the reference genome on a female hamster, we excluded reads mapping to the PAR because sequences mapping to this region could have originated from either the X or Y chromosomes and interfered with subsequent variant calling. Following mapping, we used GATK v.4.2.5.0 HaplotypeCaller (-ERC GVCF) to call SNPs then performed genotyping with genotypeGVCFs. We hard-filtered our SNPs (--mask-extension 5 “QD < 2.0” “FS > 60.0” “MQ < 40.0” “QUAL < 30.0” “DP < 10” “DP > 150”) and restricted SNPs to biallelic loci. Finally, we incorporated filtered SNPs back into the *P. sungorus* reference genome with FastaAlternateReferenceMaker to create the *P. sungorus* and *P. campbelli* pseudoreferences. For our RNASeq data, we appended query hit indexes to resulting alignment files using hisat2Tophat.py (https://github.com/goodest-goodlab/pseudo-it/tree/master/helper-scripts/hisat2Tophat.py, last accessed March 8th, 2022) to maintain compatibility with the modtools pipeline. We used our VCFs (above) to generate a mod-file for both species with vcf2mod from Lapels v.1.1.1 to convert alignments to the *P. sungorus* reference genome, and Suspenders v.0.2.6 to merge alignments while retaining the highest quality alignment per read (Holt *et al*. 2013; Huang *et al*. 2014). We used featureCounts v.2.0.1 (Liao *et al*. 2014) to estimate counts of read pairs that aligned to the same chromosome (-B and -C) and retained only singly-mapped reads. Summaries of properly mapped reads for each sample can be found in Table S2 in File S1.

We sought to compare the gene expression phenotypes observed in dwarf hamsters to those previously documented in house mice using published RNASeq data for the same four spermatogenic cell types of two subspecies of house mouse and their sterile F1 hybrids (Larson *et al*. 2017; Hunnicutt *et al*. 2022). These studies examined two subspecies of house mice, *Mus musculus musculus* (intra-subspecific F1 males between wild-derived inbred strains PWK/PhJ♀ and CZECHII/EiJ♂) and *M. m. domesticus* (intra-subspecific F1 males between wild-derived inbred strains WSB/EiJ♀ and LEWES/EiJ♂) and their sterile (PWK♀ x LEWES♂) F1 hybrids for disrupted gene expression across spermatogenesis following the same FACS protocols implemented in this study. For all comparisons between house mice and dwarf hamsters, we used read count files generated previously for house mice (Hunnicutt *et al*. 2022) and performed all subsequent analyses in parallel for both systems.

### Estimating nucleotide diversity and divergence within and between crosses and strains

We next estimated nucleotide diversity within and divergence between each strain or cross for parent species and hybrids (π and dXY). We used the bam files generated above from mapping each strain or cross to its respective reference genome (either GCA_023856395.1 for dwarf hamsters or GRCm38.p6 for house mice) to call SNPs as described above but with the addition of a step to split reads at intronic regions using the GATK function SplitNCigarReads. We generated two VCFs, one for dwarf hamster crosses and the other for house mouse crosses, which we processed and filtered separately. If an individual mouse or hamster was sequenced for more than one cell type, we randomly chose one of the represented cell types to be included in the analysis. We hard-filtered our SNPs as above and restricted SNPs to biallelic loci. Because our SNPs were called from RNASeq data and thus may be susceptible to allelic imbalances or coverage differences between samples, we next investigated how the inclusion of SNPs with different levels of missing data impacted estimation of π and dXY. We used vcftools v.0.1.17 (Danecek *et al*. 2011) to filter SNPs allowing between 0% and 90% missing data (--max_missing; Figure S3 in File S1), and filtered SNPs with a depth lower than 5 and higher than 60 (∼2-3 higher than average coverage) to eliminate multi-mapped reads. Finally, we used pixy v.1.2.10.beta2 (Korunes and Samuk 2021) on our filtered VCFs to estimate π and dXY. Patterns of nucleotide diversity across strains and crosses were qualitatively similar across missing data thresholds, so we present results corresponding to 10% missing data in the main text. However, we note that for all comparisons, estimates of π and dXY decreased with more stringent missing data thresholds regardless of strain or cross (Figure S3 in File S1).

### Gene expression pre-processing

Following read processing and mapping, we conducted all analyses in R v.4.3.1. We classified genes as “expressed” if genes had a minimum of one Fragment Per Kilobase of exon per Million mapped reads (FPKM) in at least three samples, resulting in 21,077 expressed genes across the dwarf hamster dataset and 21,212 expressed genes across the house mouse dataset. We also identified sets of genes “induced” in a given cell type defined as genes with a median expression in a given cell population (normalized FPKM) greater than two times its median expression across all other sorted cell populations (following Kousathanas *et al*. 2014). We calculated normalized FPKM values by adjusting the sum of squares to equal one using the R package vegan v.2.6-4 (Oksanen *et al*. 2013). We conducted expression analyses using edgeR v.3.42.4 (Robinson *et al*. 2010) and normalized the data using the scaling factor method (Anders and Huber 2010).

We qualitatively assessed cell population purity both by visual inspection during cell sorting and following sequencing by assessing the expression of a panel of marker genes specific to the four cell populations targeted by our FACS protocol and present in only a single copy in the *P. sungorus* annotation. Spermatogonia markers included *Dmrt1* (Raymond *et al*. 2000) and *Hells* (Green *et al*. 2018). Leptotene/zygotene markers included *Ccnb1ip1* and *Adad2* (Hermann *et al*. 2018). Diplotene was characterized by *Aurka* and *Tank* expression (Murat *et al*. 2023) and round spermatids by *Cabyr* and *Acrv1* expression (Green *et al*. 2018). To estimate relative purity, we compared mean marker gene expression across replicates for a given cell population for both parent species. A cell population was considered qualitatively pure if it had higher marker gene expression than other populations isolated by our FACS protocol and if X-linked gene expression matched the expected regulatory dynamics (*i.e.,* active vs. silenced (MSCI) vs. repressed (PSCR); Handel 2004; Namekawa *et al*. 2006). We examined expression patterns across cell populations for all genes, autosomal genes, and X-linked genes using MDS plots generated with the plotMDS function in limma v.3.56.2 (Ritchie *et al*. 2015) and heatmaps using ComplexHeatmap v.2.16.0 (Gu *et al*. 2016). MDS plots used the top 500 genes with the largest fold change difference between samples.

### Differential gene expression analysis

We assessed differential gene expression by contrasting hybrids and each parent species for all cell populations. We fit the expression data for dwarf hamsters and house mice separately with negative binomial generalized linear models with Cox-Reid tagwise dispersion estimates and adjusted P-values to a false discovery rate (FDR) of 5% (Benjamini and Hochberg 1995). We quantified the biological coefficient of variation (BCV), a metric representing the variation in gene expression among replicates (McCarthy *et al*. 2012), for each dataset. Additionally, we calculated the BCV of just parental males or hybrid samples for each species for the first three cell populations to examine whether dwarf hamster hybrids exhibited more variability in expression than house mouse hybrids and parental dwarf hamsters. For our differential expression analyses, we contrasted expression between hybrids and each parent so that a positive log fold-change (logFC) indicated overexpression in sterile males and implemented a logFC cutoff of 1.25. We then categorized differentially expressed (DE) genes into one of four categories: DE relative to only one parent species, DE relative to both parent species but with intermediate expression (intermediate), and DE relative to both parent species but outside of the range of either parent species (transgressive; Figure S2 in File S1). Unless otherwise specified, results discussed in the main text are restricted to transgressive DE genes and are presented with the logFC from the contrast of the hybrid offspring to the parent with the same X chromosome (*P. campbelli* for dwarf hamster F1 hybrids and *M. m. musculus* for house mouse F1 hybrids). Transgressive DE genes have similar logFC values regardless of which parent is used as the contrast, but figures depicting the logFC between F1 hybrids and *P. sungorus* or *M. m. domesticus* are provided in supplementary material. We also assessed differential expression between parent species/strains (parental DE) by contrasting expression between parents sharing the same X chromosome as hybrids (*i.e., M. m. musculus* and *P. campbelli*) with parents with the alternate X chromosome (*i.e., M. m. domesticus* and *P. sungorus*) so that a positive log fold-change indicated overexpression in parents with the same X chromosome as hybrids and implemented a logFC cutoff of 1.25.

We tested for significant differences in the number of under and overexpressed transgressive DE genes within a stage for both house mouse and hamster hybrids using *X*^2^ tests with chisq.tests in R and used FDR correction for multiple comparisons. We also tested for differences in the magnitude of misexpression between mouse and hamster hybrids for each stage by comparing the distributions of logFC of transgressive DE genes between hybrids and parent species using Wilcoxon signed-rank tests and FDR correction. To characterize hybrid diplotene expression in both house mice and dwarf hamsters, we used two approaches. First, we calculated Pearson’s correlation coefficient *(r)* between average normalized hybrid diplotene expression and the average normalized expression in each parental cell type. We corrected p-values for each correlation with FDR. We generated bootstrap values for each correlation coefficient by randomly sampling the expression matrices with replacement for each sample type for 1000 replicates. Second, we compared the gene sets that escaped MSCI in each species with gene sets that characterize stage-specific expression in parent species. For this analysis, we defined sets of overexpressed X-linked diplotene genes as genes with expression (normalized FPKM) in hybrids that was in the top 10% of X-linked genes in parental diplotene samples (*i.e.,* genes that normally escape MSCI). We then compared these sets of overexpressed hybrid diplotene genes to genes “induced” in each parental stage for each species. We used gProfiler2 v.0.2.3 (Kolberg *et al*. 2020) in R to perform gene ontology (GO) analysis to identify GO terms overrepresented in hybrid transgressive DE genes sets for each cell population. We only included *P. sungorus* genes associated with mouse orthologs in our GO analysis (as established by Moore *et al*. 2022), and for our background gene lists, we used *P. sungorus* genes associated with mouse orthologs that were “expressed” in hybrids and both parent species in a given stage. We retained only Biological Process GO terms with an FDR below 0.05 and ran gProfiler2 both with and without the highlight option, a two-stage algorithm for reducing resulting GO terms by grouping significant terms into sub-ontologies and then identifying the gene sets that give rise to other significant functions (Kolberg *et al*. 2020). To test whether specific chromosomes were enriched or depleted for transgressive DE genes for a given stage, we performed hypergeometric tests on the number of transgressive DE genes on a given chromosome with phyper and adjusted P-values to an FDR of 5%. We also assessed overlap in specific transgressive DE genes within stages between house mice and dwarf hamsters. We estimated whether overlap was more or less than expected by chance and whether overlapping genes were preferentially located on the X chromosome using hypergeometric tests and performed GO enrichment on overlapping genes. For all hypergeometric tests, we defined the background sets of genes as those with non-zero logFC values in both differential expression comparisons between hybrids and each parent for a given spermatogenic stage.

### Characterizing the behavior of PAR genes in dwarf hamsters

We sought to characterize the regulatory behavior of PAR genes in dwarf hamster parent species and hybrids to determine if (1) PAR genes are normally silenced in parent species (consistent with an extension of MSCI to the PAR) and (2) if PAR genes were overexpressed in hybrids (consistent with disrupted MSCI in the PAR of hybrids). The PAR on the *P. sungorus* X chromosome is on the distal arm of the X chromosome from around 115,350,000-119,112,095 bp (Moore *et al*. 2022). There are 15 annotated *P. sungorus* genes in this region (Table S3 in File S1), which is comparable to the latest PAR assembly in C57BL/6J house mice (Kasahara *et al*. 2022). Six of these are orthologous to annotated genes in mice (*Tppp2*, *Gprin1*, *Ndrg2*, *Kcnip4*, *Ndrg2*, and *Hs6st3*), but they are not located in the mouse PAR (Kasahara *et al*. 2022). Only seven of the annotated genes in the PAR were expressed in more than three replicates across all samples (Psun_G000022875, Psun_G000022880, Psun_G000022883, Psun_G000022886, *Tppp2*, *Gprin1*, and *Ndrg2*). For these genes, we assessed whether these genes were consistently expressed or silenced in parent species in any cell population and whether any genes were differentially expressed between hybrids and either species.

## RESULTS

### Impaired sperm production in hybrid male dwarf hamsters

We first established the extent of hybrid male sterility in dwarf hamsters by comparing reproductive phenotypes for *P. campbelli*, *P. sungorus*, and F1 hybrid males (Figure 1; Table S1 in File S1). *Phodopus sungorus* males had smaller testes and seminal vesicles than *P. campbelli* males (Dunn’s Test relative testes weight p < 0.001; relative seminal vesicle weight p = 0.0061; Figure 1). These differences are qualitatively consistent across independent laboratory colonies (Ishishita *et al*. 2015; Bikchurina *et al*. 2018) and likely reflect species-specific differences between *P. campbelli* and *P. sungorus*. *Phodopus sungorus* males also had lower nucleotide diversity (π = 0.00013) than *P. campbelli* males (π = 0.00045) and both house mouse parental crosses between fully inbred mouse strains (*M. m. musculus* π = 0.00029; *M. m. domesticus* π 0.00017; Figure S3 in File S1), which could contribute to depression of male fertility within this highly inbred laboratory colony (Brekke *et al*. 2018). Nucleotide divergence between parental dwarf hamster species was elevated relative to house mice (dxy = 0.0020 vs. 0.0010), consistent with reported older divergence time estimates for dwarf hamsters (Neumann *et al*. 2006). However, these dwarf hamster species did not differ in normalized sperm counts (p = 0.071) or in sperm motility (p = 0.12). The F1 hybrid males exhibited extreme reproductive defects relative to *P. campbelli* (Figure 1). Hybrid males had smaller testes (p < 0.001) and seminal vesicles (p < 0.001) than male *P. campbelli* hamsters, and importantly, produced almost no mature spermatozoa. In the one instance where a hybrid male produced a single mature spermatozoon, it was non-motile, indicating severe reproductive impairment in hybrid males. Overall, our results confirmed previous reports of reduced fertility in hybrid male dwarf hamsters (Ishishita *et al*. 2015; Bikchurina *et al*. 2018).

**Figure 1.**
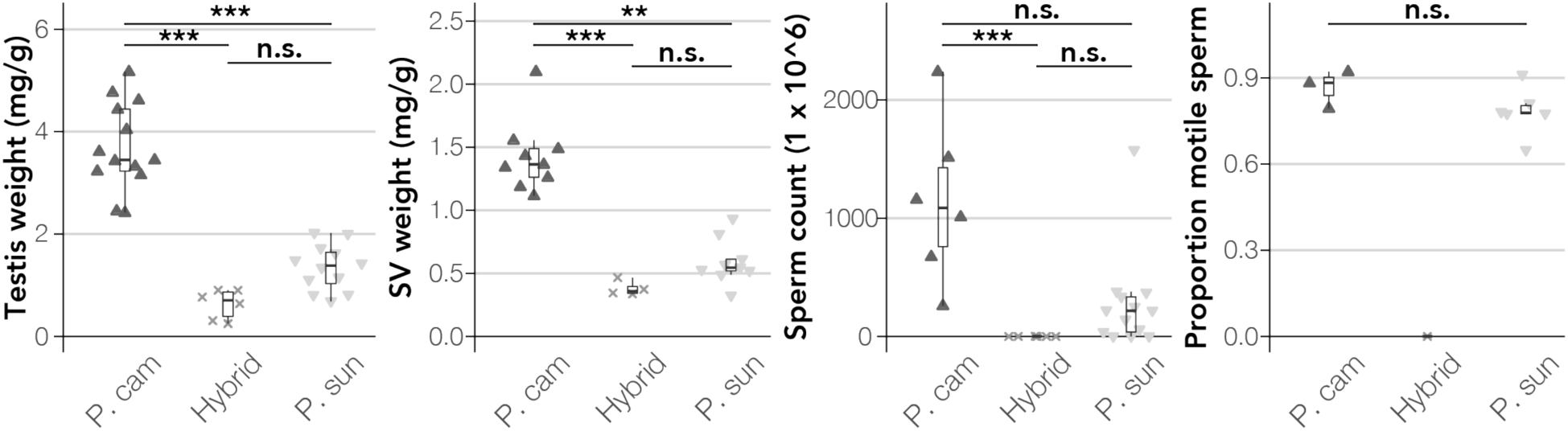
Evidence of some hybrid male sterility in dwarf hamsters. We assessed paired testes weight and paired seminal vesicle weight (SV; both normalized by body weight), sperm count, and proportion of motile sperm for *P. sungorus*, *P. campbelli*, and F1 hybrids. Whiskers extend to either the largest or smallest value or no further than 1.5 times the interquartile range, and *** indicates p < 0.001, ** indicates p < 0.01, and n.s. indicates non-significant difference between means at p > 0.05 using a post-hoc Dunn’s test with FDR correction. Upwards-pointing triangles (▴) indicate *P. campbelli*, downwards-pointing triangles (▾) indicate *P. sungorus*, and crosses (▽) indicate F1 hybrids.

### Cell type-specific gene expression across spermatogenesis

To characterize cell type-specific gene expression, we used FACS to isolate enriched cell populations from each fertile parent species and their sterile F1 hybrids across four stages of spermatogenesis. The four targeted populations included: spermatogonia (mitotic precursor cells), leptotene/zygotene spermatocytes (meiotic cells before MSCI), diplotene spermatocytes (meiotic cells after MSCI), and round spermatids (postmeiotic cells). We were unable to isolate round spermatids from the F1 hybrids, which was consistent with the lack of mature spermatozoa present in the cauda epididymis extractions (Figure 1). We sequenced RNA from each cell population for *P. campbelli* (spermatogonia n = 4, leptotene/zygotene n = 5, diplotene n = 5, and round spermatids n = 4; Table S2 in File S1), *P. sungorus* (spermatogonia n = 4, leptotene/zygotene n = 4, diplotene n = 5, and round spermatids n = 3), and F1 hybrid males (*P. campbelli*♀ x *P. sungorus*♂; spermatogonia n = 4, leptotene/zygotene n = 4, diplotene n = 4). We compared our hamster expression data to an analogous cell type-specific RNASeq dataset from two species of house mice, *Mus musculus musculus* and *M. m. domesticus*, and their sterile F1 hybrids (n = 3 for all cell populations in each cross; Larson *et al*. 2017).

We used two approaches to qualitatively evaluate the purity of spermatogonia, leptotene/zygotene spermatocytes, diplotene spermatocytes, and round spermatids isolated from males from both parental species: (1) we quantified the relative expression of a panel of cell population marker genes, and (2) we characterized the expression patterns of the sex chromosomes across development. We found that our candidate marker genes had the highest expression in their expected cell population for all stages except leptotene/zygotene (Figures S4 and S5 in File S1), indicating high purity of spermatogonia, diplotene spermatocytes, and round spermatids. Leptotene/zygotene markers did not have the highest expression in leptotene/zygotene samples (except for *Ccnb1ip1*), potentially indicating lower purity of this cell population. Nonetheless, the patterns of X chromosome expression in fertile parents were consistent with expectations across this developmental timeline: the X chromosome had active expression in spermatogonia and leptotene/zygotene cells, was inactivated in diplotene cells consistent with MSCI, and was partially inactivated in round spermatids, consistent with PSCR (Figure 2; Namekawa *et al*. 2006), indicating successful isolation of these cell populations.

**Figure 2.**
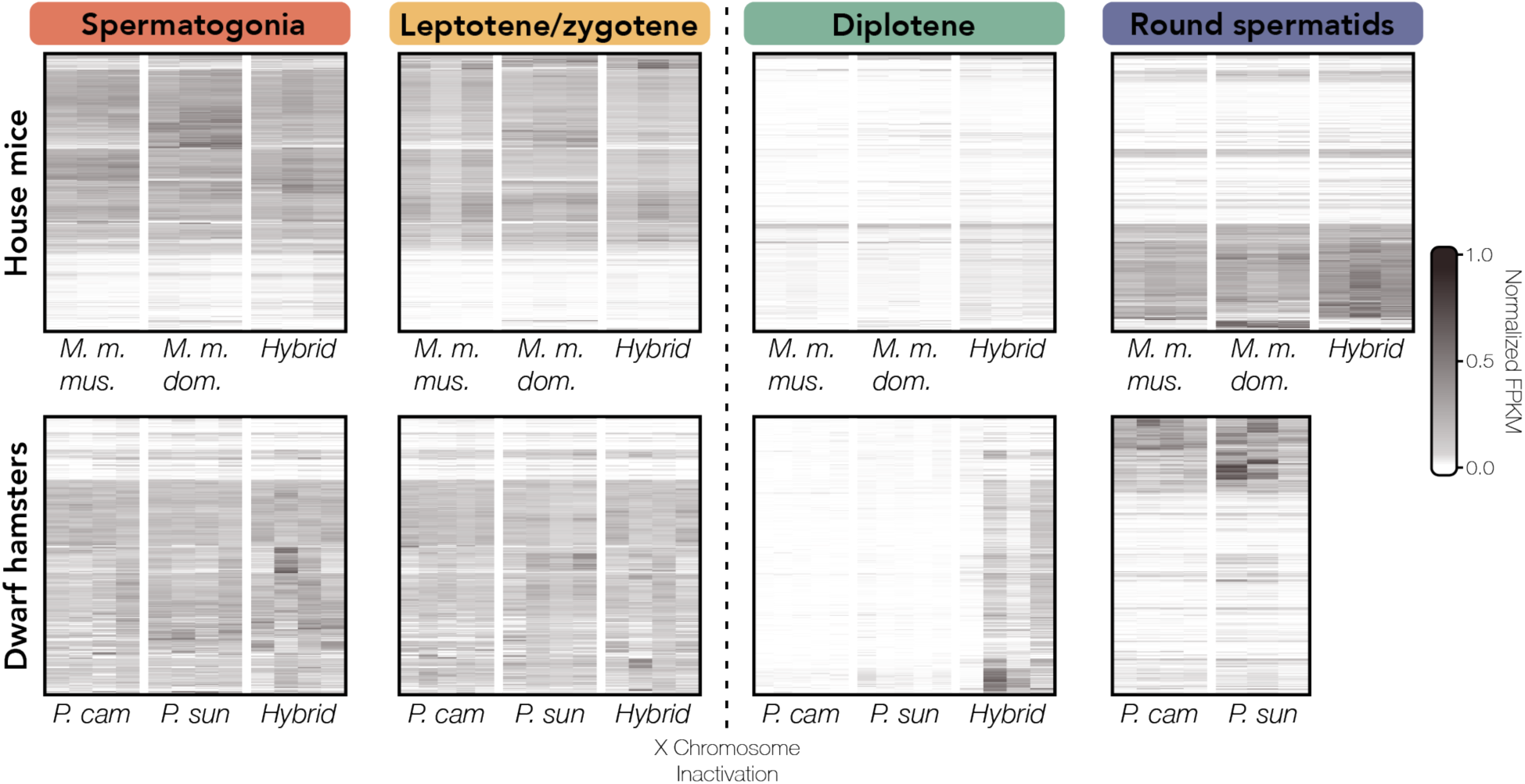
Overexpression of X-linked genes in diplotene spermatocytes in both house mouse and dwarf hamster hybrids. Heatmap of X-linked gene expression in house mice (upper panel) and dwarf hamsters (lower panel) plotted as normalized FPKM values that are hierarchically clustered using Euclidean distance. Each column represents a different individual, each row represents a gene, and darker colors indicate higher expression. The heatmap was generated with the R package ComplexHeatmap v.2.12.0 (Gu et al. 2016). Note, hybrid dwarf hamsters do not produce mature spermatozoa, and accordingly, we were unable to isolate round spermatids.

When we examined overall expression differences within dwarf hamsters and within house mice, we found that samples clustered primarily by cell population on MDS1 and 2 (Figure 3), then by cross/strain when cell populations were examined separately (Figures S6 and S7 in File S1). Within all cell populations across both systems, hybrids showed intermediate overall expression patterns to parent species (Figures S6 and S7 in File S1). However, in dwarf hamsters but not house mice, spermatogonia and leptotene/zygotene samples overlap rather than forming distinct clusters (Figure 3). Further, the expression profiles of hybrid diplotene cell populations ranged from clustering with parental leptotene/zygotene to parental diplotene cell populations, which contrasted with what we observed in house mice where hybrid diplotene cell populations clustered distinctly with parental diplotene cell populations (Figure 3). Overall, our results indicate that we successfully isolated cell populations in dwarf hamsters that span key stages of spermatogenesis.

**Figure 3.**
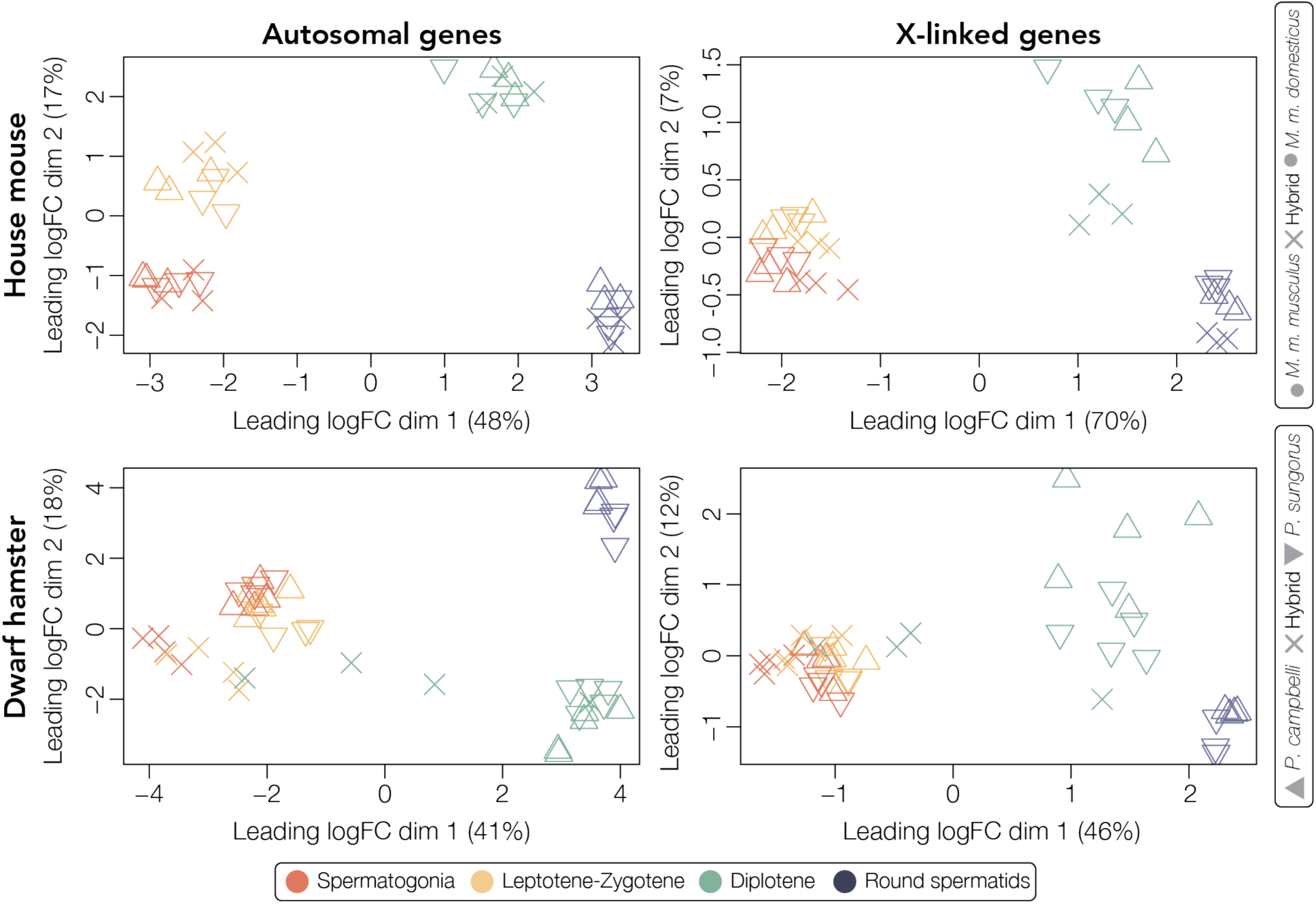
Hybrid gene expression profiles cluster by parental spermatogenic cell population in house mice but not dwarf hamsters. Multidimensional scaling (MDS) plots of distances among house mouse (upper panels) and dwarf hamster (lower panels) samples for expressed autosomal (left) and X-linked (right) genes. Distances are calculated as the root-mean-square deviation (Euclidean distance) of log2 fold changes among the top 500 genes that distinguish each sample. Each strain or cross is indicated by a symbol, and samples are colored by cell population.

### Disrupted transcription early in spermatogenesis in dwarf hamsters

We sought to characterize which expression phenotypes were associated with sterile hybrids in both house mice and dwarf hamsters. We first investigated whether differential gene expression in hybrids tended towards intermediate or transgressive expression, and then within transgressive DE genes, whether misexpression tends towards up- or downregulation in hybrids compared to parents. Differential gene expression in mouse hybrids had a slight bias towards transgressive expression except in round spermatids (percentage of transgressive DE genes: SP: 72.0%, LZ: 62.7%, DIP: 60.8%, and RS: 24.9%; Figures 4b and S2 in File S1), while almost all differential expression in dwarf hamster hybrids was transgressive (percentage of transgressive DE genes: SP: 99.2%, LZ: 98.0%, and DIP: 97.1). For transgressive DE genes, autosomal misexpression in hybrid house mice was biased towards upregulation across spermatogenesis (mean logFC of spermatogonia autosomal transgressive DE genes: +1.90/ *X*^2^: p < 0.001; leptotene/zygotene: +1.33/ *X*^2^: p < 0.001; diplotene: +0.47/ *X*^2^: p < 0.001; Figures 4c and S8 in File S1), as was X-linked misexpression (spermatogonia mean logFC: +1.98 / *X*^2^: p < 0.001; leptotene/zygotene: +2.50/ *X*^2^: p < 0.001; diplotene: +2.78/ *X*^2^: p < 0.001). In contrast, we found that the direction of misexpression in dwarf hamster hybrids was cell type-specific. Autosomal transgressive DE genes in dwarf hamster hybrids were overwhelmingly downregulated in both early stages of spermatogenesis, especially in comparison to house mice (mean logFC of spermatogonia autosomal transgressive DE genes: -4.38/ *X*^2^: p < 0.001; leptotene/zygotene: -4.67/ *X*^2^: p < 0.001; Figure 4d) but upregulated in diplotene (average logFC = +4.67/ *X*^2^: p < 0.001; Figures 4d and S8 in File S1). When comparing both X-linked and autosomal expression in dwarf hamsters, we found similar patterns: almost all X-linked transgressive DE genes in the first two stages of spermatogenesis were exclusively downregulated (spermatogonia mean logFC = -6.57/ *X*^2^: p < 0.001; leptotene/zygotene: -6.73/ *X*^2^: p < 0.001), but misexpression was biased towards upregulation in diplotene (mean logFC: 5.70/ *X*^2^: p < 0.001).

**Figure 4.**
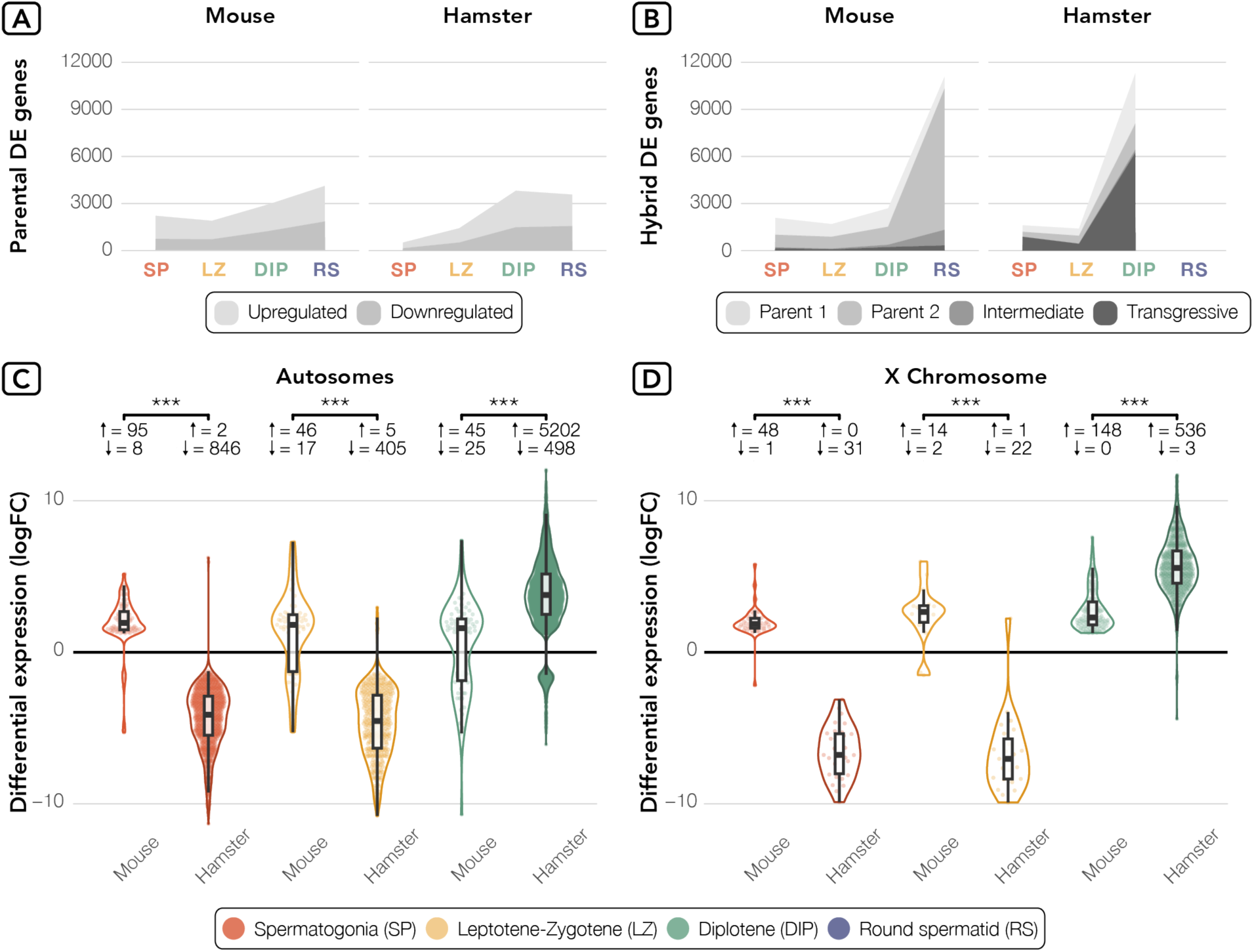
House mice and dwarf hamster hybrids have opposite patterns of disrupted regulation early in spermatogenesis. a) Up- and down-regulated DE genes between parent species for house mice (left; *M. m. musculus* vs. *M. m. domesticus*) and dwarf hamsters (right; *P. sungorus* vs. *P. campbelli*). b) Counts of DE genes between hybrids and one parent species (two lightest shades of gray where Parent 1 was either *M. m. musculus or P. sungorus* and parent 2 was either *M. m. domesticus* or *P. campbelli)* or between hybrids and both parents (two darker shades of gray) that showed either intermediate or transgressive expression. c) Transgressive DE genes in house mouse and dwarf hamster hybrids for autosomal genes where the logFC represents hybrid expression relative to *M. m. musculus* or *P. campbelli,* respectively, and d) transgressive DE gene expression for X-linked genes. Results are displayed for autosomes (left) and the X chromosome (right). *** indicates p < 0.001 for pairwise comparisons from Wilcoxon signed-rank tests after FDR correction. Whiskers extend to either the largest or smallest value or no further than 1.5 times the interquartile range. The number of up and downregulated transgressive DE genes in hybrids are listed next to arrows indicating direction of differential expression.

Second, we investigated how developmental stage influenced the extent of hybrid misexpression. In both systems, the number of DE genes between parent species increased with the progression of spermatogenesis, consistent with less constraint on gene expression levels as spermatogenesis progresses (Figure 4a; Kopania *et al*. 2022a; Murat *et al*. 2023). Similarly, transgressive misexpression in both hybrid dwarf hamsters and house mice increased with the progression of spermatogenesis, though to a greater extent in dwarf hamsters (Figure 4b). While both parental DE genes and transgressive hybrid DE genes increased with the progression of spermatogenesis, parental DE genes initially exceeded the number of transgressive hybrid DE genes in hybrids in spermatogonia and leptotene/zygotene spermatocytes. However, in diplotene spermatocytes, transgressive hybrid differential expression surpassed parental differential expression in both house mice and dwarf hamsters. We also found a much greater genome-wide disruption of expression in diplotene cell populations of hybrid dwarf hamsters than in hybrid house mice (Figures 4c, 4d, and S8-S10 in File S1), indicating more widespread regulatory disruption. We then compared transgressive DE genes between house mice and dwarf hamster hybrids across all three stages and found that the number of shared transgressive DE genes did not differ from the number expected by chance in early spermatogenesis (spermatogonia = 3 genes, hypergeometric test p = 0.98; leptotene/zygotene = 2 genes, p = 0.59; Table S4 in File S1). However, the number of shared transgressive DE genes did exceed the number expected by chance in diplotene cell populations (n = 68; p < 0.001), and these shared genes were preferentially located on the X chromosome (61/68; p < 0.001). Additionally, these sets of shared transgressive DE genes between house mice and dwarf hamsters were not significantly enriched for any GO terms, though many play known roles in spermatogenesis, the apoptotic process, and cell differentiation (Table S5 in File S1) and may be promising candidates for future functional analysis. Ultimately, we found that there are similar trends in the patterns of transgressive expression across stages in both systems, and although few specific genes had disrupted expression in both dwarf hamster and house mouse hybrids, genes with similar patterns of disrupted expression tend to be X-linked and disrupted in later stages of spermatogenesis.

To determine if the differences we observed in the extent of misexpression between dwarf hamsters and house mice was due to greater expression variability across our dwarf hamster samples, we calculated the BCV, a measurement of inter-replicate variability, for each species and hybrid across the first three cell populations. Inter-replicate variability was higher in dwarf hamster hybrids relative to house mouse hybrids (dwarf hamster BCV = 0.69; house mice = 0.18). Additionally, nucleotide diversity within dwarf hamster hybrids was higher (π = 0.0012) than within house mouse hybrids (π = 0.00052; Figure S3 in File S1). However, the extent of the expression variability observed in hybrids relative to the inter-replicate variability of parental species differed between house mice and dwarf hamsters: dwarf hamster hybrid variability was more than dwarf hamster parental samples (*P. campbelli* = 0.49; *P. sungorus* = 0.40), but hybrid variability was similar to parental samples for house mice (*M. m. musculus* = 0.19; *M. m. domesticus* = 0.22). The greater inter-replicate expression variability in dwarf hamster hybrids relative to parent species suggests that the increased misexpression we see in hybrids cannot be explained by greater inter-replicate variability in our dwarf hamster samples alone, and it may also reflect the greater nucleotide diversity present among dwarf hamster hybrids.

Third, we characterized whether there was a clear difference between the autosomes and sex chromosomes in expression phenotype by testing whether the X chromosome was enriched for transgressive DE genes in each stage for both systems. Across all stages of spermatogenesis in house mice, the X chromosome was enriched for transgressive DE genes between hybrids and parents (spermatogonia p < 0.001; leptotene/zygotene p < 0.001; diplotene p < 0.001; round spermatids p < 0.0036; Figure S9 in File S1). In contrast, misexpression in dwarf hamsters was not uniformly sex chromosome-specific across all stages, as the sex chromosomes in dwarf hamsters showed no enrichment for transgressive DE genes early in spermatogenesis (spermatogonia: p = 0.21; Figure S10 in File S1) despite X chromosome enrichment in both leptotene/zygotene (p = 0.0064) and round spermatids (p < 0.001). However, we note that the magnitude of misexpression was greater for sex chromosomes than autosomes in dwarf hamsters across all stages (Figures 4c and 4d; discussed above). Only two autosomes were enriched for transgressive DE genes in dwarf hamsters in any cell population: one scaffold on chromosome 5 in spermatogonia (JAJQIY010003390.1; hypergeometric test; p = 0.003) and chromosome 11 in leptotene/zygotene (p = 0.017; Figure S10 in File S1). Together, the subtle differences in the distribution of transgressive DE genes across autosomes and the X chromosome between dwarf hamster hybrids and house mouse hybrids suggest a difference in the extent of the role for sex chromosome-specific disruption between systems.

### Misexpression in diplotene appears to be unrelated to disrupted MSCI in dwarf hamsters

We next asked whether expression patterns indicated similar disrupted regulatory processes resulting in sex chromosome-specific misexpression in both systems. In sterile hybrid house mice, the mean logFC of transgressive X-linked DE genes during diplotene was higher than the mean logFC of transgressive autosomal DE genes (X logFC = +2.78 in contrast to autosomal logFC = +0.47; Figures 4c and 4d), consistent with disrupted MSCI (Good *et al*. 2010; Bhattacharyya *et al*. 2013; Campbell *et al*. 2013; Turner and Harr 2014; Larson *et al*. 2017, 2022). In sterile dwarf hamster hybrids, we also found elevated mean logFC of transgressive X-linked DE genes relative to autosomal genes (X logFC = +5.70 in contrast to autosomal logFC = +3.69), but the extent of X chromosome overexpression, as measured by logFC of X-linked transgressive DE genes, was greater than in hybrid house mice (5.7/3.69 or ∼1.5x higher; Figures 2, 4c, and 4d). There was also more variability in the extent of overexpression of X-linked genes in hybrid dwarf hamsters compared to normal parental X-linked expression during diplotene (relative overexpression = 18.5 +/-6.8) than for overexpression of X-linked genes in hybrid house mice compared to normal parental X-linked expression in diplotene (relative overexpression = 1.75 +/- 0.09; Figures 2 and S11 in File S1). Strikingly, some hybrid male dwarf hamsters had an almost completely silenced X chromosome, while others had an almost completely transcriptionally-activated X chromosome (Figure 2).

Despite overexpression of the X chromosome during diplotene in hybrid dwarf hamsters, the overall expression phenotype, including the identity and the extent of misexpression of overexpressed genes, appeared to fundamentally differ between house mouse and dwarf hamster hybrids (Figures 2 and 5a-5c). We established these differences in X-linked overexpression using two approaches. First, we tested which parental cell types had the highest expression correlation with hybrid diplotene cell types for both X-linked and autosomal genes. In mice, the expression profile of X-linked genes in hybrids during diplotene was most positively correlated with the expression profile of X-linked parental round spermatid genes, consistent with disrupted MSCI (spermatogonia *(r)* = -0.25, p < 0.001; leptotene/zygotene *(r)* = -0.073, p = 0.035; round spermatid *(r)* = 0.26, p < 0.001; Figures 2 and 5a). Autosomal diplotene genes showed no positive correlations with either spermatogonia (*r* = -0.26; p < 0.001), leptotene/zygotene (*r* = -0.032; p < 0.001), or round spermatids (*r* = -0.060; p < 0.001; Figure 5a). If the X-linked overexpression phenotype in dwarf hamsters was consistent with disrupted MSCI, then we would also expect X-linked and autosomal expression profiles in hybrid diplotene to follow the same patterns. In contrast to this prediction, hybrid dwarf hamster diplotene expression profiles for both X-linked and autosomal genes had a positive correlation with parental leptotene/zygotene (autosomal *(r)* = 0.14, p < 0.001; X-linked *(r)* = 0.22, p < 0.001) and spermatogonia (autosomal *(r)* = 0.027, p < 0.001; X-linked *(r)* = 0.091, p = 0.016) and a negative correlation with round spermatids (autosomal *(r)* = -0.30, p < 0.001; X-linked *(r)* = -0.31, p < 0.001; Figure 5b). These striking differences in the strength and direction of expression profile correlations suggest that the regulatory mechanisms underlying the overexpression phenotype of X-linked genes in sterile hybrids differed between house mice and dwarf hamsters.

**Figure 5.**
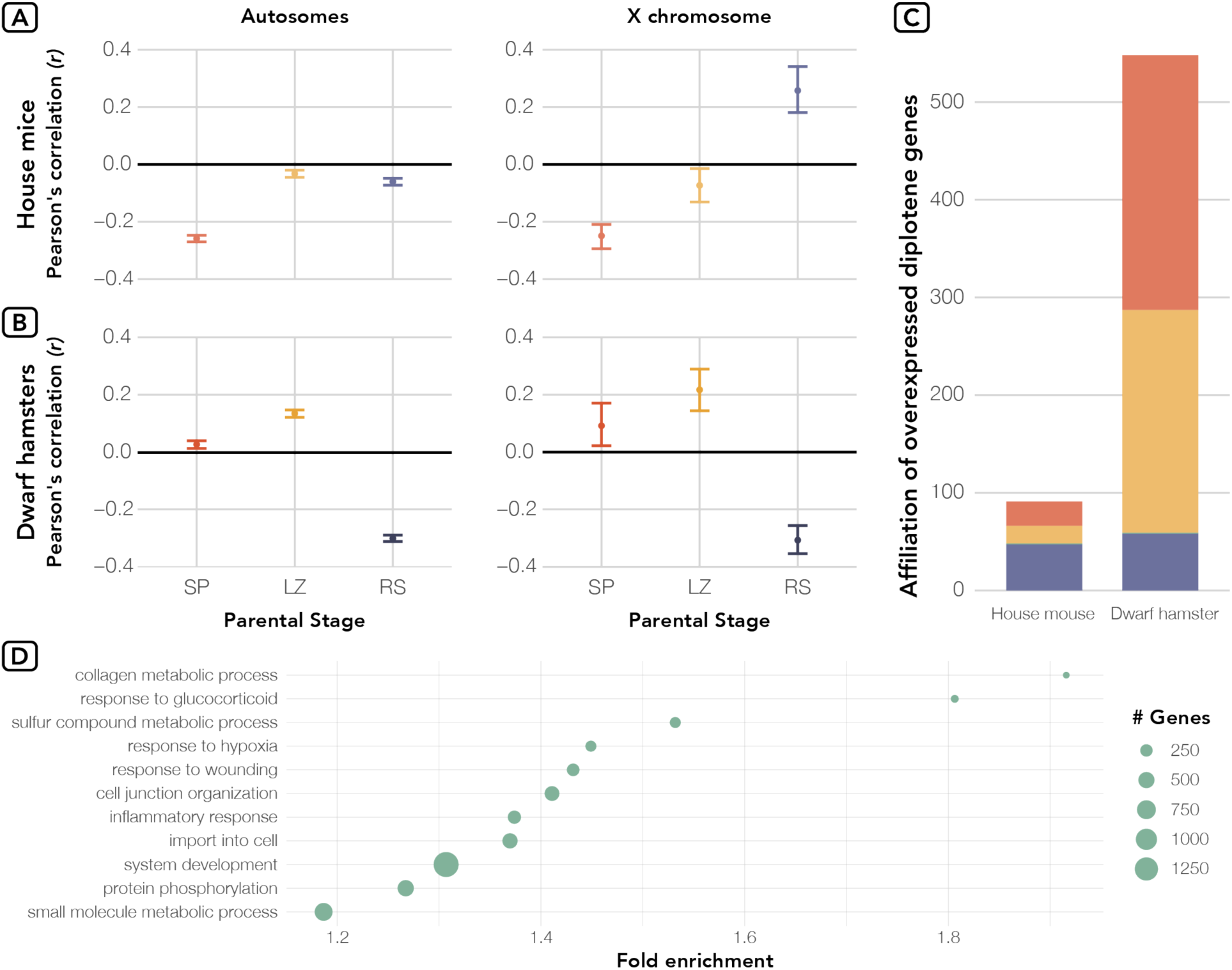
Spermatogenesis in hybrid dwarf hamsters appears to stall after leptotene/ zygotene. a) We calculated the Pearson’s correlation coefficient *(r)* between mean hybrid diplotene expression and the mean expression for each parental cell type for both house mice and b) dwarf hamsters. Correlation coefficients were calculated for both autosomal (left panels) and X-linked (right panels) genes. We then generated bootstrap values by randomly sampling the expression matrices for 1000 replicates. All correlation coefficients were significantly different from zero (p < 0.05) after FDR correction. c) Classification of X-linked overexpressed genes in diplotene cell populations of hybrid mice and dwarf hamsters by parental stage in which genes are induced. Genes are colored by cell type (red = spermatogonia, yellow = leptotene/zygotene, green = diplotene, and blue = round spermatids). d) Select enriched Biological Process GO terms (ranked by FDR) for transgressive DE genes between hybrid dwarf hamsters and *P. campbelli* in diplotene spermatocytes. Included terms shown are the result of the highlight function within gProfiler2 which collapses GO terms in a two-step clustering algorithm (Kolberg *et al*. 2020). Point size corresponds to the number of genes belonging to each GO term, and terms are plotted by the fold enrichment of the GO term in the dataset relative to the provided gene backgrounds.

This difference in pattern was further supported when we compared which sets of genes were overexpressed in hybrid diplotene in dwarf hamsters and house mice. For this approach, we looked at parental gene expression patterns to characterize which stages of spermatogenesis all genes were normally active during and characteristic of (*i.e.,* “induced”; see Methods). Using this information, we then identified which X-linked genes were overexpressed in hybrid diplotene (defined as genes with normalized expression in the top 10% of X-linked genes) and assessed which parental stages the overexpressed X-linked genes were characteristic of in both systems. As in our correlation analysis, we found that in hybrid house mice, the genes that were overexpressed in diplotene most closely resembled genes that are normally active in round spermatids in parental mice (51.6% of genes), but that in hybrid dwarf hamsters, overexpressed diplotene genes resembled spermatogonia- and leptotene/zygotene-specific genes (47.6% and 41.6% respectively; Figure 5c). Because of the dissimilarity in the genes that were overexpressed during diplotene in both systems, we next performed gene ontology (GO) enrichment analyses on the set of transgressive DE genes in hybrids to determine which biological processes could be potentially contributing to this pattern. In contrast to house mice hybrids where transgressive DE genes were enriched for no biological processes, the transgressive DE genes in dwarf hamster were enriched for several biological processes including cell junction/extracellular matrix organization, system development, inflammatory response, and apoptotic process which together point to widespread disruption of numerous processes necessary for normal male fertility (highlighted terms presented in Figure 5d and the full lists in Tables S6-S8 in File S1). Collectively, these results suggest that the X-linked overexpression phenotype in sterile hybrid dwarf hamsters is inconsistent with disrupted MSCI and is possibly related to a stalling and breakdown of spermatogenesis between leptotene/zygotene and diplotene during Prophase I.

### PAR expression was not disrupted in sterile hybrid dwarf hamsters

Finally, we tested the hypothesis that hybrid sterility in dwarf hamsters may be correlated with asynapsis of the X and Y chromosomes because of divergence in the pseudoautosomal region (PAR) which prevents proper chromosome pairing (Bikchurina *et al*. 2018). The PAR is the only portion of the sex chromosomes that is able to synapse during routine spermatogenesis, and PAR genes are assumed to escape silencing by MSCI (Raudsepp and Chowdhary 2015). However, because XY asynapsis is common in dwarf hamster hybrids, Bikchurina et al. (2018) hypothesized that MSCI may extend to the PAR of hybrid male dwarf hamsters, resulting in the silencing of PAR genes in hybrids that may be critical to meiosis (Figure 6a). To test this hypothesis, we compared the expression of genes located in the dwarf hamster PAR (Moore et al. 2022) between parental dwarf hamster species and hybrid offspring across the timeline of spermatogenesis. Specifically, we hypothesized that if XY asynapsis results in an extension of MSCI to the PAR in hybrid dwarf hamsters, then hybrids should have similar PAR gene expression to parents early in meiosis before homologous chromosome synapse during pachytene. This should be followed by the silencing of PAR genes in hybrids, but not parent species, during diplotene. If XY asynapsis does not alter the regulation of the PAR in hybrids, then we may see two possible patterns. First, if all PAR genes are critical to the later stages of spermatogenesis, then PAR genes in both hybrids and parents should be uniformly expressed in diplotene. Alternatively, if PAR genes are not critical to the later stages of spermatogenesis, then hybrids and parent species should have similar PAR gene expression, and not all PAR genes may be expressed during diplotene.

**Figure 6.**
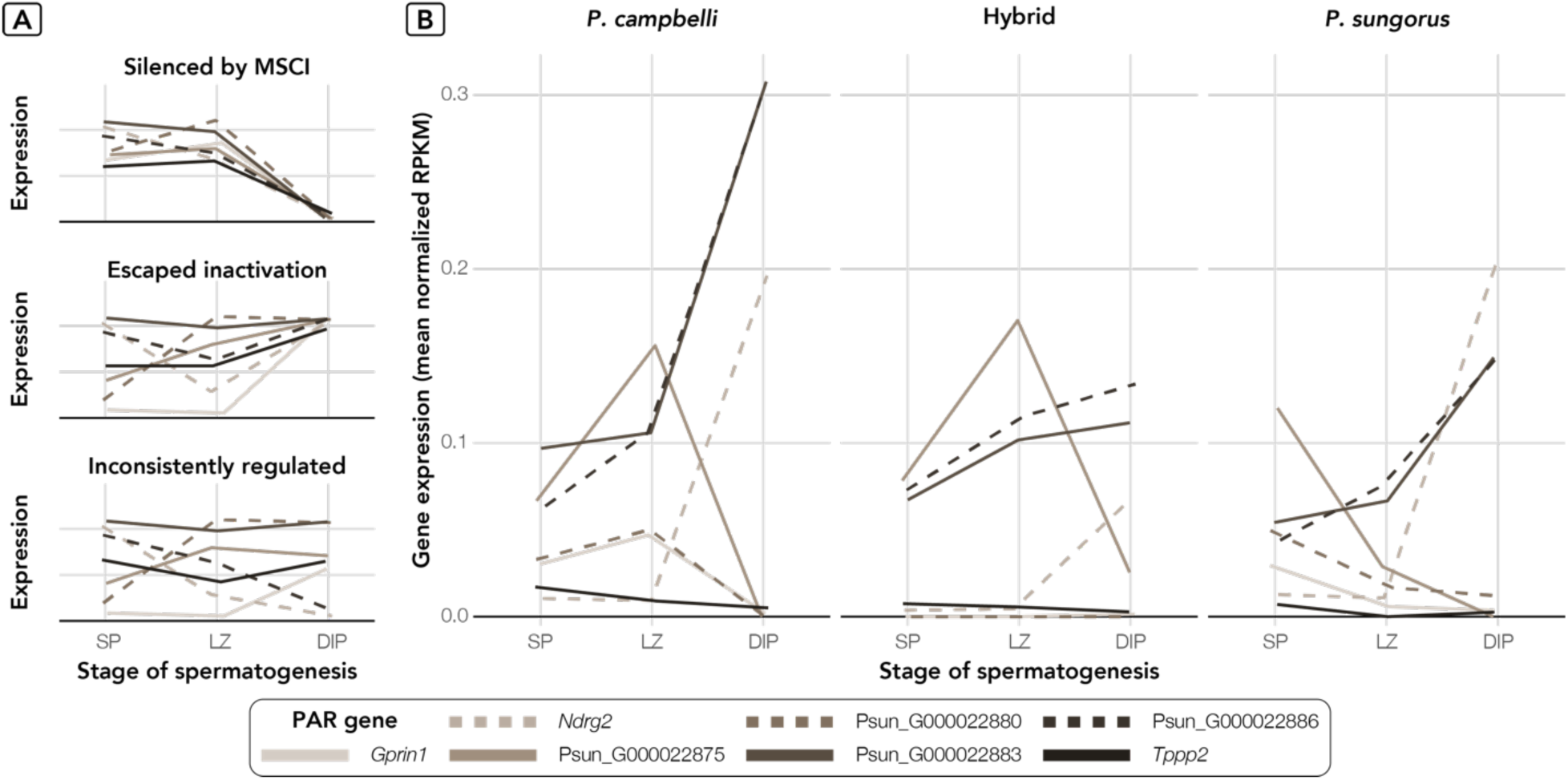
PAR gene expression is not disrupted during spermatogenesis in hybrid dwarf hamsters. a) Hypothesized types of PAR gene expression across spermatogenesis. If MSCI extends to the entire X chromosome, then PAR genes would show some level of expression early in spermatogenesis which would then drop to zero in diplotene when MSCI occurs. If the PAR escapes silencing by MSCI and if PAR genes are critical to spermatogenesis, then we would expect PAR genes to be uniformly expressed in diplotene when the rest of the X chromosome is silenced. Finally, if the PAR escapes silencing by MSCI but all PAR genes are not critical to spermatogenesis, then we would expect some PAR genes to be expressed and some to be not expressed in diplotene. b) Observed patterns of PAR gene expression (as mean normalized RPKM across individuals) in parental species and hybrid dwarf hamsters for all PAR genes (indicated by line type and color) across spermatogenesis (SP = spermatogonia, LZ = leptotene/zygotene., and DIP = diplotene).

We did not find evidence supporting PAR-wide silencing in dwarf hamster hybrids during diplotene suggesting that MSCI is not extended to the PAR in dwarf hamster hybrids because of XY asynapsis (Figure 6b). Furthermore, we do not see PAR-wide expression of genes during diplotene in hybrids or parents, indicating that not all PAR genes are critical to the progression of spermatogenesis in dwarf hamsters. In general, most PAR gene expression followed similar trends between hybrids and parent species. Two PAR genes were differentially expressed between hybrids and *P. campbelli* during diplotene (*Ndrg2* and Psun_G000022883; Table S3 in File S1), but these genes were still expressed in hybrids. Further, an association between PAR misregulation during the early stages of spermatogenesis and hybrid male sterility also seems unlikely as only one gene, *Gprin1*, showed transgressive differential expression in early meiosis between both parent species and hybrids (Table S3 in File S1). Thus, based on the current annotation of the PAR in *P. sungorus*, we currently find no direct evidence linking improper silencing of PAR genes to hybrid male sterility in dwarf hamsters.

## DISCUSSION

We used a comparative approach to understand common gene expression phenotypes associated with hybrid male sterility in two divergent rodent crosses. We characterized the asymmetry in the expression patterns of transgressive genes, how misexpression changed over developmental timelines, and how the X chromosome and autosomes differed in both of these aspects. We found that while there were similarities in hybrid expression phenotypes in house mice and dwarf hamsters, there were also differences in the timing and chromosomal distribution of disrupted gene expression that point towards different underlying mechanisms behind hybrid male sterility.

### Asymmetry and developmental timing of misexpression in hybrids

We first investigated patterns of transgressive gene misexpression in sterile male hybrids. Studies of transgressive misexpression in sterile or inviable hybrids have often focused on whether hybrid expression is biased towards over or underexpression, with the hypothesis that expression may be biased towards overexpression if hybrid incompatibilities disrupt repressive gene regulatory elements (Meiklejohn *et al*. 2014; Barreto *et al*. 2015; Larson *et al*. 2017). In house mice, there is strong support for overexpression of both autosomal and X-linked genes in sterile F1 hybrids (Mack *et al*. 2016; Larson *et al*. 2017, 2022; Hunnicutt *et al*. 2022). Surprisingly, we found that in dwarf hamster hybrids, there was nearly uniform downregulation of transgressive DE genes in mitotic and early meiotic cell populations, suggesting that a loss of regulatory repression is not an inevitable outcome of hybrid genomes. Hybrid house mice expression is also more similar to the parent with the same X chromosome, *M. m. musculus,* than to the parent with a different X chromosome, *M. m. domesticus* (Figure 4b and S2 in File S1; Larson *et al*. 2017). Further work in house mice has shown that F1 hybrid expression patterns depend on both autosomal background and sex chromosome mismatch (Kopania *et al*. 2022b). In contrast, hybrid dwarf hamsters showed similar levels of misexpression in both the *P. campbelli* and *P. sungorus* comparisons. Determining what factors shape the misexpression of parental alleles is a fruitful area of future research.

Asymmetric patterns of misexpression have been found in many hybrids, including underexpression in sterile *Drosophila* hybrids (Michalak and Noor 2003; Haerty and Singh 2006; Llopart 2012) and sterile introgression lines of tomato (Guerrero *et al*. 2016) and *Drosophila* (Meiklejohn *et al*. 2014), but overexpression has also been found in other sterile hybrids (Llopart 2012; Davis *et al*. 2015). In many of these studies, patterns of misexpression may be complicated by differences in cell composition or differences in the developmental timeline of sterile hybrids and their parents (Good *et al*. 2010; Wei *et al*. 2014; Montgomery and Mank 2016; Kerwin and Sweigart 2020; Hunnicutt *et al*. 2022). The variation we and others have found in hybrid expression phenotypes suggests that the mechanisms of disrupted expression are complex, even within groups with relatively shallow divergence times, such as rodents, and we need more data from diverse hybrid sterility systems to begin to understand common drivers of transgressive hybrid misexpression.

The downregulation we observed in early spermatogenesis in hybrid dwarf hamsters could be due to impaired transcription factor binding with promoter or enhancer elements (Oka *et al*. 2014; Guerrero *et al*. 2016) or disrupted epigenetic silencing. Disruption of epigenetic regulation of gene expression has been increasingly linked to hybrid dysfunction in plants (Shivaprasad *et al*. 2012; Lafon-Placette and Köhler 2015; Zhu *et al*. 2017), especially polyploids (Paun *et al*. 2007), and may also contribute to hybrid male sterility in *Drosophila* (Bayes and Malik 2009) and cattle x yak hybrids (Luo *et al*. 2022). At least one known chromatin difference, an expansion of the heterochromatin-enriched Xp arm of the X chromosome, has been documented between parental dwarf hamster species (Gamperl *et al*. 1977; Haaf *et al*. 1987). However, it is unknown what the functional consequences of this chromatin state divergence or other diverged epigenetic regulatory mechanisms, such as methylation, may be in hybrid dwarf hamster spermatogenesis, and further work is needed to distinguish between potential mechanisms underlying the observed genome-wide downregulation.

Spermatogenesis as a developmental process may be sensitive to disruption (Lifschytz and Lindsley 1972; Wu and Davis 1993), but it remains an open question whether specific stages of spermatogenesis, or developmental processes more broadly, may be more prone to the accumulation of hybrid incompatibilities. In general, earlier developmental stages are thought to be under greater pleiotropic constraint and less prone to disruption (Cutter and Bundus 2020). With the progression of mouse spermatogenesis, pleiotropy decreases (as approximated by increases in tissue specificity; Murat *et al*. 2023) and the rate of protein-coding evolution increases (Larson *et al*. 2016; Kopania *et al*. 2022a; Murat *et al*. 2023), which may make the later stages of spermatogenesis more prone to accumulating hybrid incompatibilities. Indeed, we found fewer DE genes both between parent species and in sterile hybrids for both dwarf hamsters and house mice during the early stages of spermatogenesis than in later stages (Figures S9 and S10), and hybrid misexpression greatly exceeds parental expression divergence in late spermatogenesis. When examining transgressive DE genes shared between analogous cell types in dwarf hamster and house mouse hybrids, we found that there were similar or fewer shared genes than expected by chance during early spermatogenesis but more shared genes than expected by chance, especially on the X chromosome, in later spermatogenesis, suggesting that disrupted expression of shared genes of large effect during early spermatogenesis is unlikely to be responsible for the repeated evolution of hybrid male sterility in these species.

Despite general similarities in patterns of misexpression across spermatogenesis in hybrids, studies in house mice suggest that early spermatogenesis may be tolerant of some misregulation as low levels of gene misexpression in early meiotic stages does not always correlate with a complete cessation of sperm development (Oka *et al*. 2010; Ishishita *et al*. 2015; Mipam *et al*. 2023). However, the patterns we find in dwarf hamsters suggests that spermatogenesis may be disrupted between zygotene and diplotene cell stages from early misexpression. Dwarf hamster hybrid diplotene cell populations had X-linked and autosomal gene expression profiles which more closely resemble parental leptotene/zygotene cell populations than either diplotene or postmeiotic cell populations. Furthermore, transgressive DE genes in hybrids during diplotene were enriched for genes associated with cell differentiation, proliferation, and programmed cell death, suggesting misexpression during this stage could be a consequence of a stalling or breakdown of early meiosis in hybrid dwarf hamsters. It’s unclear what underlying genomic mechanisms could result in this breakdown, but it is possible that this disruption could potentially act as a major contributor to hybrid sterility in this system. Ultimately, we find that spermatogenesis is a complex and rapidly evolving developmental program that may provide many potential avenues across its timeline for the evolution of hybrid incompatibilities.

### Abnormal sex chromosome expression patterns differ between dwarf hamster and house mouse hybrids

The sex chromosomes play a central role in speciation, an observation which has been supported by both Haldane’s rule (Haldane 1922) and the large X-effect on hybrid male sterility (Coyne and Orr 1989). Misregulation of the X chromosome may contribute to hybrid sterility in several species pairs (Davis *et al*. 2015; Morgan *et al*. 2020; Sánchez-Ramírez *et al*. 2021). The X chromosome is transcriptionally repressed during routine spermatogenesis in many organisms including eutherian mammals (McKee and Handel 1993), monotremes (Murat *et al*. 2023), *Drosophila* (Landeen *et al*. 2016), grasshoppers (Viera *et al*. 2021), mosquitos (Taxiarchi *et al*. 2019), and nematodes (Rappaport *et al*. 2021). Because of the ubiquity of X chromosome repression during spermatogenesis, disruption of transcriptional repression could be a widespread regulatory phenotype in sterile hybrids (Lifschytz and Lindsley 1972; Larson *et al*. 2018). In sterile hybrid mice, disrupted X repression (disrupted MSCI) leads to the overexpression of the normally silenced X chromosome during diplotene (Good *et al*. 2010; Bhattacharyya *et al*. 2013; Campbell *et al*. 2013; Turner and Harr 2014; Larson *et al*. 2017, 2022). We also found overexpression of the X chromosome during diplotene in sterile hybrid dwarf hamsters, but in a manner inconsistent with sterile hybrid house mice. Both the X chromosome and autosomes are overexpressed in dwarf hamster hybrid diplotene cell populations to a greater extent on average than was observed in house mice, and importantly, X-linked overexpression was more variable in dwarf hamster hybrids than house mice hybrids. In fact, some dwarf hamster hybrids had wildly overexpressed X chromosomes while others appeared to have properly silenced X chromosomes. Our expression correlation and gene set analyses of hybrid diplotene cell populations provide additional evidence that the genes overexpressed in hybrid hamster diplotene are different than those overexpressed in house mouse hybrids, sharing more similarity to the earlier meiotic cell types than downstream postmeiotic cell types. Overall, our results indicate fundamentally different patterns of X-linked overexpression in both systems, with X-linked overexpression in dwarf hamster hybrids being inconsistent with disrupted MSCI patterns observed in house mouse hybrids.

Much of what we know about the genomic architecture and the role of sex chromosome misregulation in hybrid male sterility in mammals comes from decades of work that have shown a major gene, *Prdm9*, and its X chromosome modulator, *Hstx2*, may be responsible for most F1 hybrid male sterility in house mice (Forejt *et al*. 1991, 2021; Trachtulec *et al*. 1997; Mihola *et al*. 2009; Lustyk *et al*. 2019). *Prdm9* directs the location of double strand breaks during meiotic recombination (Mihola *et al*. 2009; Oliver *et al*. 2009; Smagulova *et al*. 2016). In hybrid mice, divergence at *Prdm9* binding sites leads to asymmetric double-stranded breaks and results in autosomal asynapsis, triggering Meiotic Silencing of Unsynapsed Chromatin, shutting down transcription on asynapsed autosomes using the same cellular machinery as MSCI (Turner 2015), and eventually meiotic arrest and cell death (Bhattacharyya *et al*. 2013; Forejt *et al*. 2021). This process is associated with the disruption of MSCI and a characteristic overexpression of the X chromosome during meiosis (Good *et al*. 2010; Bhattacharyya *et al*. 2013; Campbell *et al*. 2013; Turner and Harr 2014; Larson *et al*. 2017, 2022), but whether disrupted MSCI directly contributes to hybrid male sterility or is simply a downstream consequence of *Prdm9* divergence is still uncertain (Forejt *et al*. 2021).

Whether we should have expected patterns of disrupted sex chromosome expression in sterile hybrid hamsters to be the same as house mice is unclear. The sex chromosomes in pachytene cells of hybrid dwarf hamsters display normal yH2AFX staining (Ishishita *et al*. 2015; Bikchurina *et al*. 2018), a key marker in MSCI (Abe *et al*. 2022), which may indicate that the hybrid sex chromosomes are properly silenced. Additionally, autosomal asynapsis is rarely observed in hybrid dwarf hamsters, and asynapsis is almost exclusive to the sex chromosomes (Ishishita *et al*. 2015; Bikchurina *et al*. 2018). This contrasts *Prdm9*-mediated sterility in house mice, where hybrid autosomes are often asynapsed and decorated with yH2AFX (Bhattacharyya *et al*. 2013; Forejt *et al*. 2021). Mechanisms other than *Prdm9* may also disrupt MSCI and result in sterility, such as macrosatellite copy number divergence (Bredemeyer *et al*. 2021) and X-autosome translocations that impair synapsis (Homolka *et al*. 2007), although there is no evidence for X-autosome translocations between these two species of dwarf hamsters (Moore *et al*. 2022). Thus, while MSCI may be a major target for the accumulation of reproductive barriers between species in many mammalian systems, either through *Prdm9* divergence or alternative mechanisms, our results suggest that sterility in dwarf hamsters has a more composite regulatory basis.

Another mechanism often proposed to underlie mammalian male hybrid sterility, especially in rodents, is divergence in the PAR between parental species. The PAR is the only portion of the sex chromosomes which can synapse during spermatogenesis, and it is still unclear if the regulation of the PAR is uniformly detached from MSCI across divergent mammalian sex chromosome systems (Raudsepp and Chowdhary 2015). We find no evidence that PAR-specific misregulation is associated with hybrid sterility in dwarf hamsters. PAR genes are not silenced in hybrids or parents, a pattern that is inconsistent with MSCI that has extended to the PAR due to XY asynapsis, and further, expression of PAR genes in hybrids differs little from parental PAR expression. While we find no evidence that PAR misregulation *per se* is associated with hybrid sterility in this system, we cannot rule out the possibility that structural and sequence divergence between the PARs of *P. sungorus* and *P. campbelli* may be associated with hybrid sterility. Structural and sequence divergence in the PAR has been hypothesized to activate the meiotic spindle checkpoint by interfering with proper pairing of sex chromosomes (Burgoyne *et al*. 2009; Dumont 2017). The PAR evolves rapidly in rodents (White *et al*. 2012b; Raudsepp and Chowdhary 2015; Morgan *et al*. 2019), and this elevated divergence may underlie sex chromosome asynapsis and apoptosis in several hybrid mouse crosses (Matsuda *et al*. 1991; Oka *et al*. 2010; White *et al*. 2012a; Dumont 2017). Furthermore, divergence in the mouse PAR has been implicated in spermatogenic defects in crosses where *Prdm9*-divergence is minimal, such as between closely related subspecies (Dumont 2017) or in mice with genetically-modified *Prdm9* alleles (Davies *et al*. 2021). Meiosis is likely tolerant to some degree of divergence in the PAR (Morgan *et al*. 2019), but exact limits are currently unknown. At this time, thorough analysis of structural and sequence divergence between the PARs in dwarf hamsters is challenging as the PAR is notoriously difficult to assemble (but see Kasahara *et al*. 2022), and there are annotation gaps in the current assembly of the PAR in dwarf hamsters. In sum, we find no clear pattern of regulatory disruption of PAR genes in sterile hybrid dwarf hamsters, though this result may change pending further refinement of the PAR annotation.

### Conclusions

Cell-specific approaches for quantifying expression phenotypes are powerful tools for providing insight into the underlying mechanisms behind hybrid dysfunction (Hunnicutt *et al*. 2022), especially in systems where it remains difficult to interrogate the underlying genomic architecture of these traits. Using a contrast of dwarf hamster and house mouse hybrids, we have shown that transgressive overexpression is not an inevitable outcome of hybridization, that misexpression resulting from hybrid incompatibilities may be likely to arise in differing stages of spermatogenesis, and that disrupted sex chromosome silencing does not appear to play an equal role in sterility between these two systems. Both the expression phenotypes we observed here and histological evidence from other studies (Ishishita *et al*. 2015; Bikchurina *et al*. 2018) suggest that several reproductive barriers are acting during spermatogenesis in dwarf hamster hybrids. It has become increasingly apparent as more study systems are investigated that the genetic basis of postzygotic species barriers are often complex and polymorphic (Cutter 2012; Coughlan and Matute 2020), and implementing approaches which account for the developmental complexities of hybrid dysfunction, as we have done here, will allow us to make further advances in understanding the processes of speciation.

## Acknowledgements

This work was supported by a National Science Foundation Graduate Research Fellowship to KEH (DGE-2034612) and the following grants awarded to KEH: Sigma Xi Grant-in-aid of Research, American Society of Mammalogists Grant-in-Aid of Research Grant, and a Society for the Study of Evolution R.C. Lewontin Early Award. Additionally, this work was supported by grants from the Eunice Kennedy Shriver National Institute of Child Health and Human Development of the National Institutes of Health (R01-HD073439, R01-HD094787) to JMG, and the National Science Foundation to ELL (DEB-2012041). We would like to thank members of the Larson and Good labs, as well as Jonathan Velotta and members of the Velotta lab, for feedback on this project, Gabrielle Welsh for technical assistance, Ivan Kovanda for support using the University of Denver Research Data Analysis Cluster, and Pamela K. Shaw and the UM Fluorescence Cytometry Core supported by an Institutional Development Award from the NIGMS (P30GM103338). Any opinions, findings, and conclusions or recommendations expressed in this material are those of the author(s) and do not necessarily reflect the views of the National Science Foundation or the National Institutes of Health.

## Author contributions

KEH, JMG, and ELL conceived of the study. CC, SK, ELL, and KEH conducted lab work. KEH conducted the analyses. KEH and ELL wrote the manuscript with input from ECM, CC, SK, and JMG.

## Data accessibility

Hamster sequence data are available through the NCBI SRA under accession number PRJNA1024468, and mouse sequence data are available under SRA accession PRJNA296926. File S1 contains all supplemental figures. Code used for the analyses and fertility phenotypic data are available from GitHub (https://github.com/KelsieHunnicutt/dwarf_hamster_hybrids).

