## Supplementary material for "Different complex regulatory phenotypes underlie hybrid male sterility in divergent rodent crosses": File S1

**Supplemental Figures:**

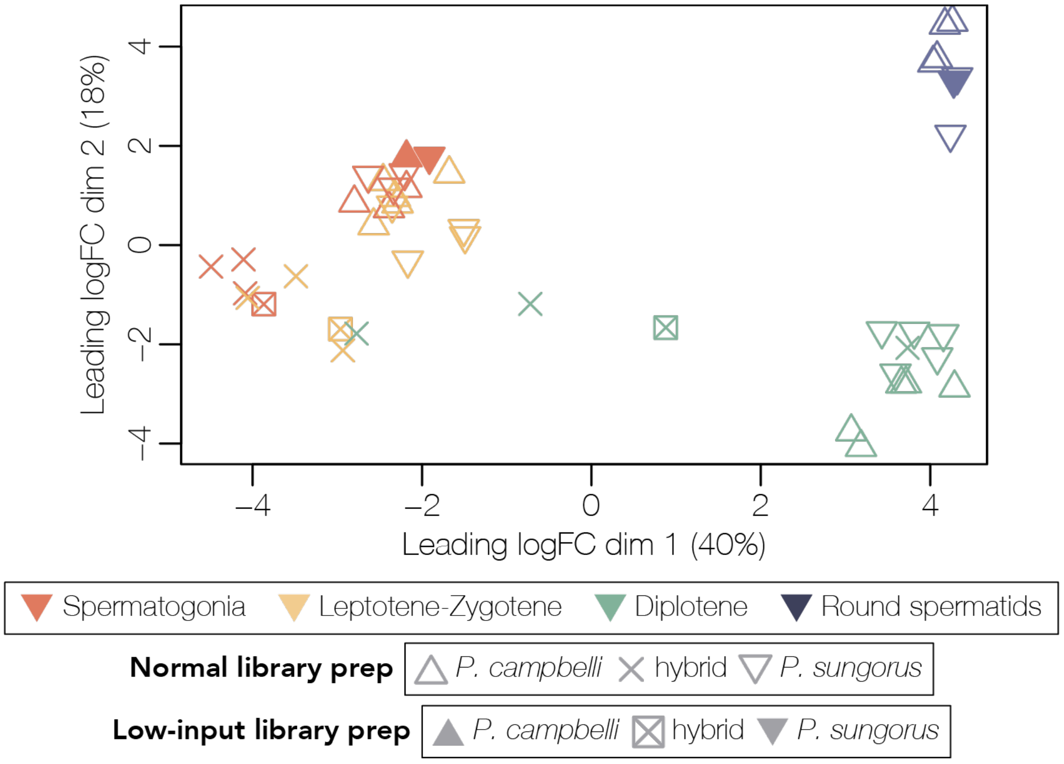


**Figure S1. Library preparation had minimal batch effects on samples.** Multidimensional scaling (MDS) plots of distances among dwarf hamster samples (lower panels). Distances are calculated as the root-mean-square deviation (Euclidean distance) of log2 fold changes among genes that distinguish each sample. Library type as well as each species and F1 hybrid are indicated by symbols, and samples are colored by cell population (red = spermatogonia, yellow = leptotene/zygotene, green = diplotene, blue = round spermatids).

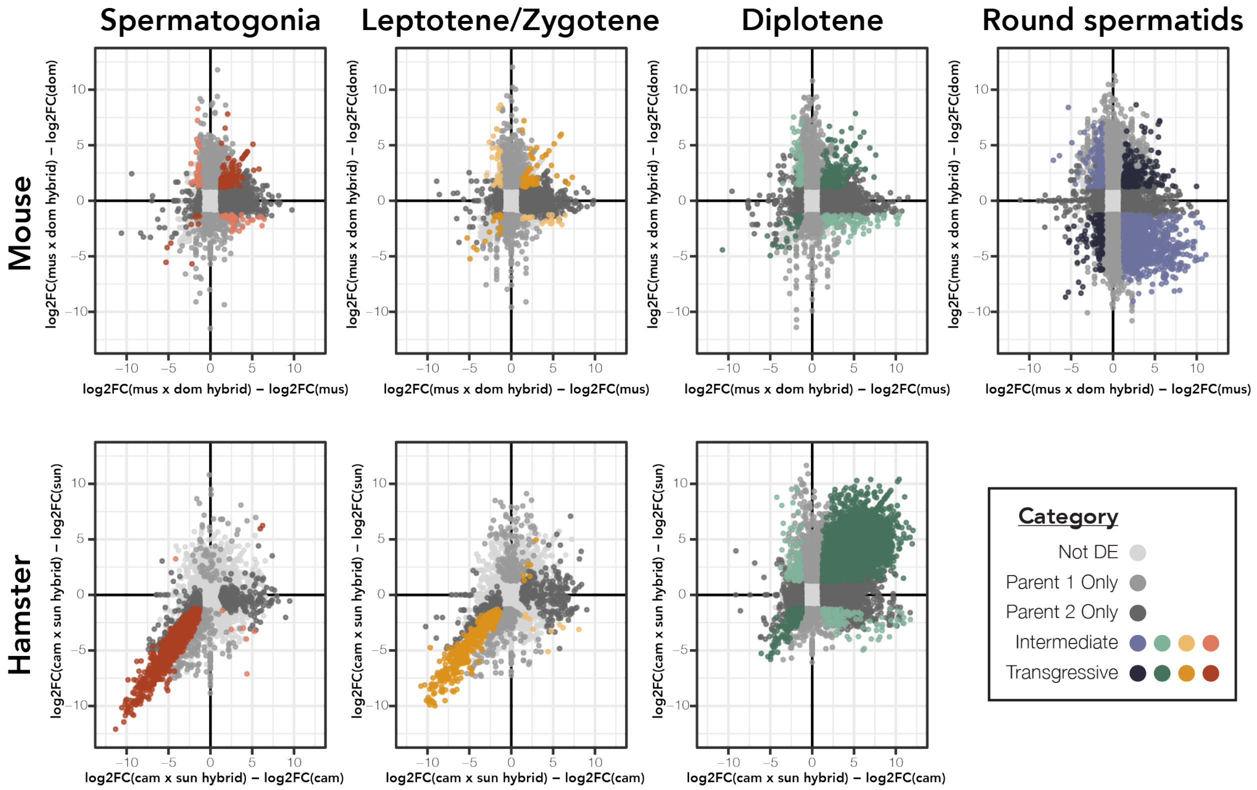


**Figure S2. Most DE genes between hybrids and both parents exhibit transgressive expression patterns regardless of system or stage of spermatogenesis.** For each stage of spermatogenesis, we identified DE genes between house mouse or dwarf hamster hybrids and either only one or both of their respective parent species and required a logFC greater than 1.5. Genes that were DE only with respect to one parent are in the two darker shades of gray: Parent 1 was either *M. m. musculus* or *P. sungorus* and Parent 2 was either *M. m. domesticus* or *P. campbelli*. Genes that were DE between hybrids and both parents were then categorized as either intermediate in expression (less saturated colors) or (outside of the range of either parent; more saturated colors).

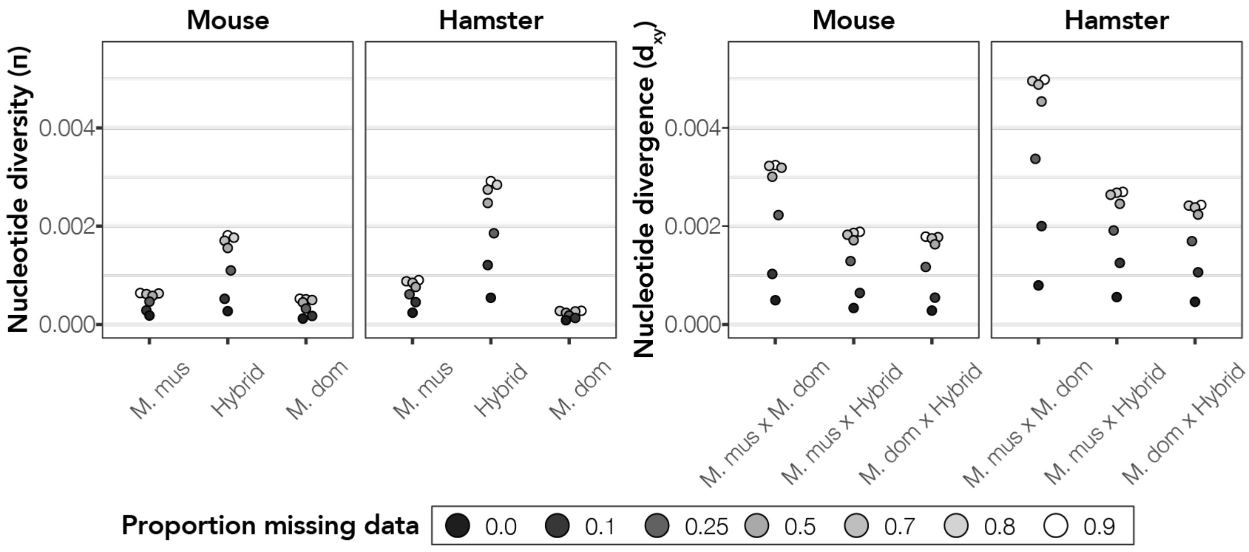


**Figure S3. Nucleotide diversity is lowest within *P. sungorus* parental individuals consistent with some inbreeding depression in this laboratory colony.** We estimated nucleotide diversity within (π; left panel) and between house mice and dwarf hamster strains/crosses ($d_{XY}$; right panel) using pixy (Korunes and Samuk 2021) across different amounts of permitted missing data (indicated by color saturation).

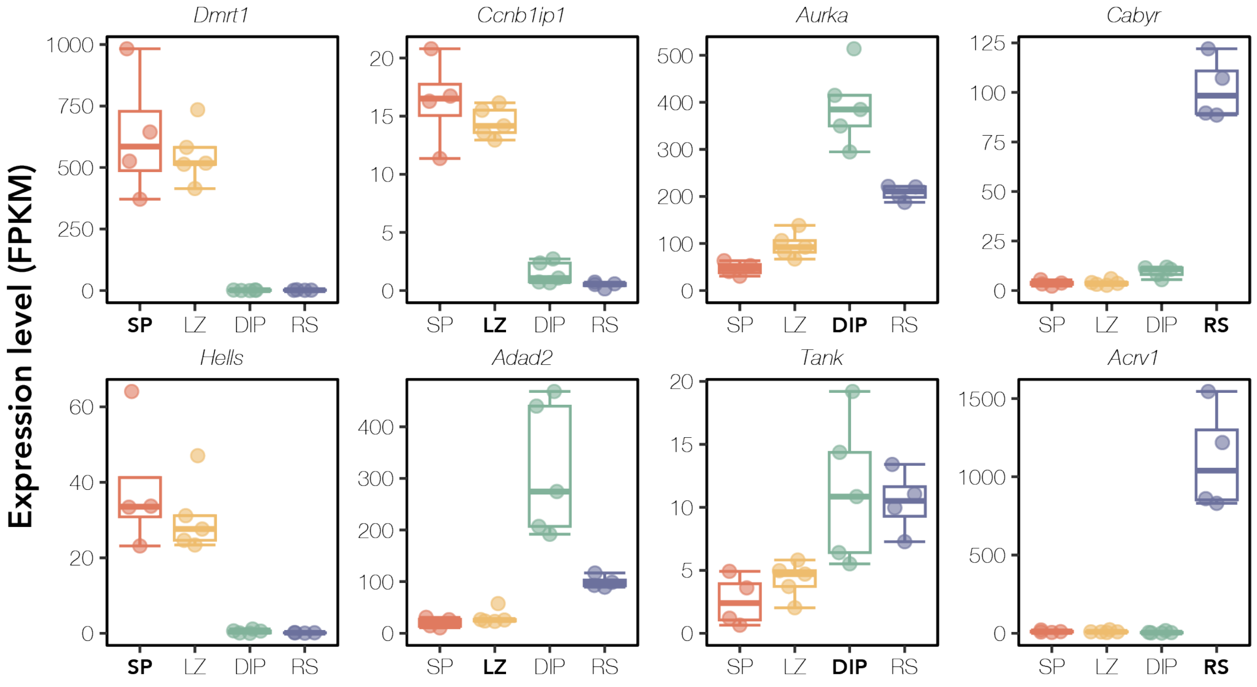


**Figure S4. Marker genes indicate successful FACS of target cell populations across spermatogenesis for *P. campbelli* samples.** Expression of cell type-specific marker genes across each sample type for *P. campbelli* reference samples. We quantified expression (FPKM) of two marker genes (rows) associated with testes-specific cell types (columns). Each panel displays marker expression in each sample type (red = spermatogonia, yellow = leptotene/zygotene, green = diplotene, blue = round spermatids). Sample types are bolded in each panel where marker gene expression is expected. *Dmrt1* (Raymond *et al.* 2000) and *Hells* (Green *et al.* 2018) are spermatogonia markers, *Ccnb1ip1* and *Adad2* (Hermann *et al.* 2018) are leptotene/zygotene markers, *Aurka* and *Tank* (Murat *et al.* 2023) are diplotene markers, and *Cabyr* and *Acrv1* (Green *et al.* 2018) are round spermatid markers. Note, that while *Adad2* has low expression in leptotene/zygotene samples, *P. campbelli* leptotene/zygotene samples have expected X chromosome expression (Figure 4; main text).

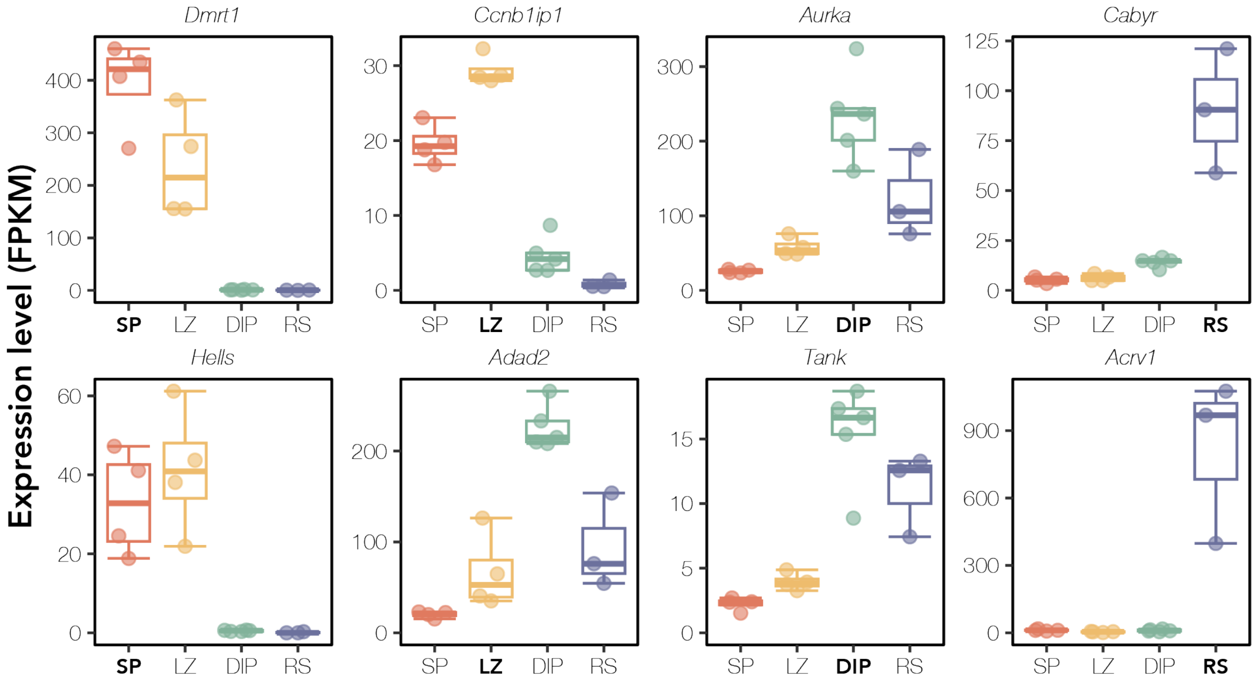


**Figure S5. Marker genes indicate successful FACS of target cell populations across spermatogenesis for *P. sungorus* samples.** Expression of cell type-specific marker genes across each sample type for *P. sungorus* reference samples. We quantified expression (FPKM) of two marker genes (rows) associated with testes-specific cell types (columns). Each panel displays marker expression in each sample type (red = spermatogonia, yellow = leptotene/zygotene, green = diplotene, blue = round spermatids). Sample types are bolded in each panel where marker gene expression is expected. *Dmrt1* (Raymond *et al.* 2000) and *Hells* (Green *et al.* 2018) are spermatogonia markers, *Ccnb1ip1* and *Adad2* (Hermann *et al.* 2018) are leptotene/zygotene markers, *Aurka* and *Tank* (Murat *et al.* 2023) are diplotene markers, and *Cabyr* and *Acrv1* (Green *et al.* 2018) are round spermatid markers. Note, that while *Adad2* has low expression in leptotene/zygotene samples, *P. sungorus* leptotene/zygotene samples have expected X chromosome expression (Figure 4; main text).

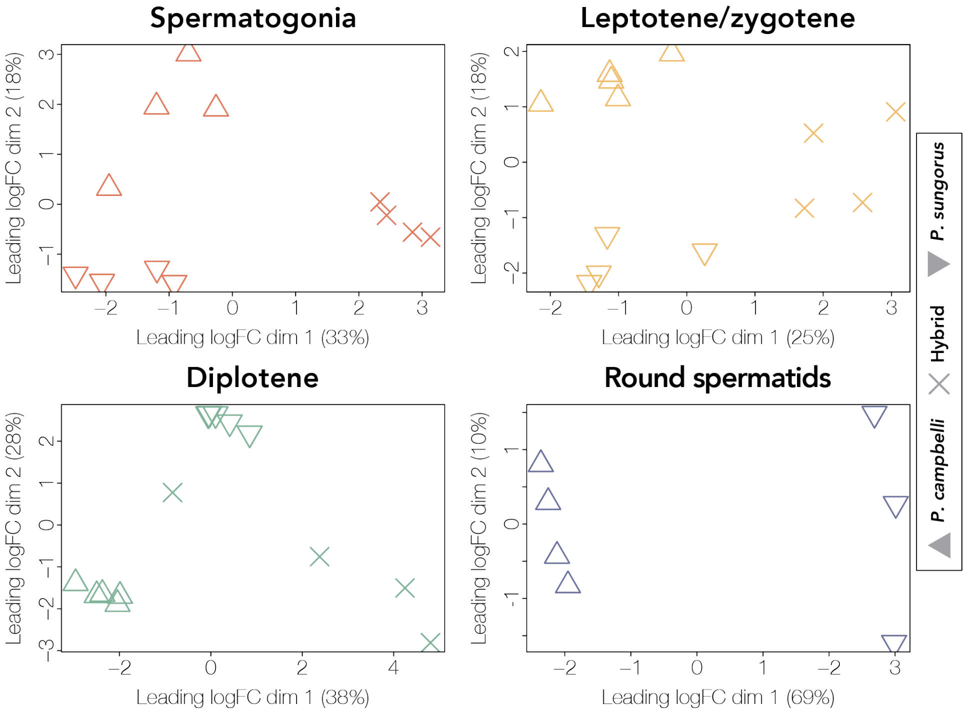


**Figure S6. Hybrid dwarf hamsters have intermediate expression to parent species within each cell population.** Multidimensional scaling (MDS) plots of distances among dwarf hamster samples for each cell population. Distances are calculated as the root-mean-square deviation (Euclidean distance) of log2 fold changes among genes that distinguish each sample. Each cross or strain is indicated by a symbol (*P. campbelli* = △, *P. sungorus* = ▽, and hybrid = ✖). Samples are colored by cell population (red = spermatogonia, yellow = leptotene/zygotene, green = diplotene, blue = round spermatids).

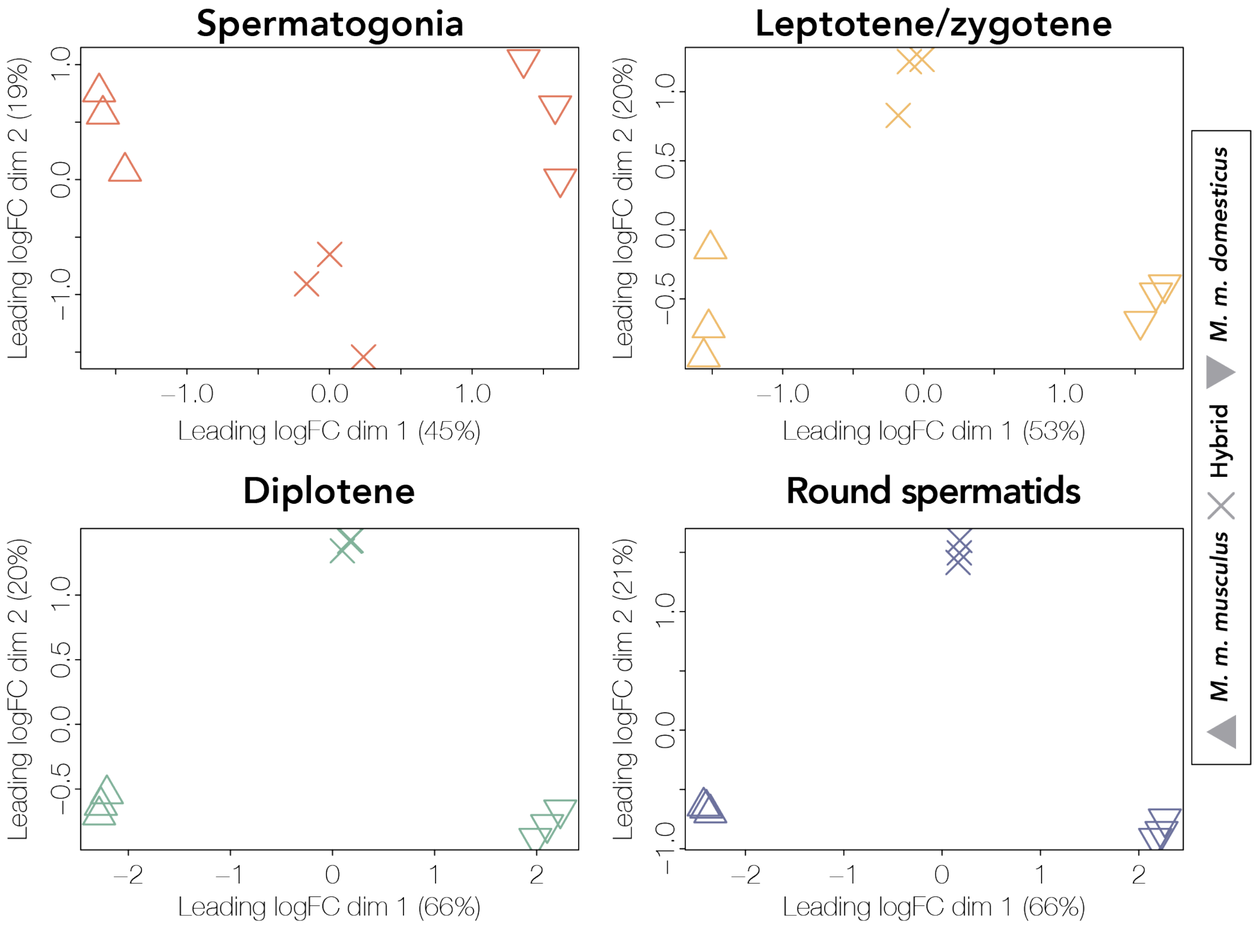


**Figure S7. Hybrid house mice have intermediate expression to parent species within each cell population.** Multidimensional scaling (MDS) plots of distances among house mouse samples for each cell population. Distances are calculated as the root-mean-square deviation (Euclidean distance) of log2 fold changes among genes that distinguish each sample. Each cross or strain is indicated by a symbol (*M. m. musculus* = △, *M. m. domesticus* = ▽, and sterile hybrid = ✖). Samples are colored by cell population (red = spermatogonia, yellow = leptotene/zygotene, green = diplotene, blue = round spermatids).

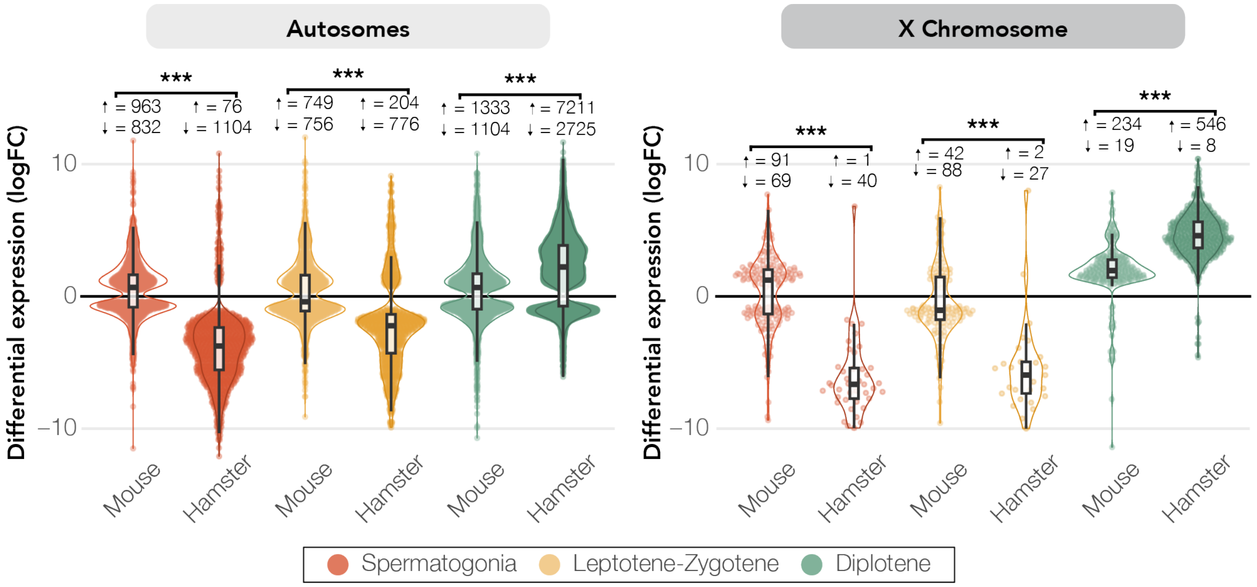


**Figure S8. Opposite patterns of disrupted regulation early in spermatogenesis between house mice and dwarf hamsters.** Transgressive DE genes in house mouse and dwarf hamster hybrids for autosomal genes (left), where the logFC represents hybrid expression relative to *M. m. domesticus* or *P. sungorus*, respectively, and transgressive DE gene expression for X-linked genes (right). Results are displayed for autosomes (left) and the X chromosome (right). *** indicates $p < 0.001$ for pairwise comparisons from Wilcoxon signed-rank tests after FDR correction. Whiskers extend to either the largest or smallest value or no further than 1.5 times the interquartile range. The number of up and downregulated DE genes in hybrids are listed next to arrows indicating direction of differential expression.

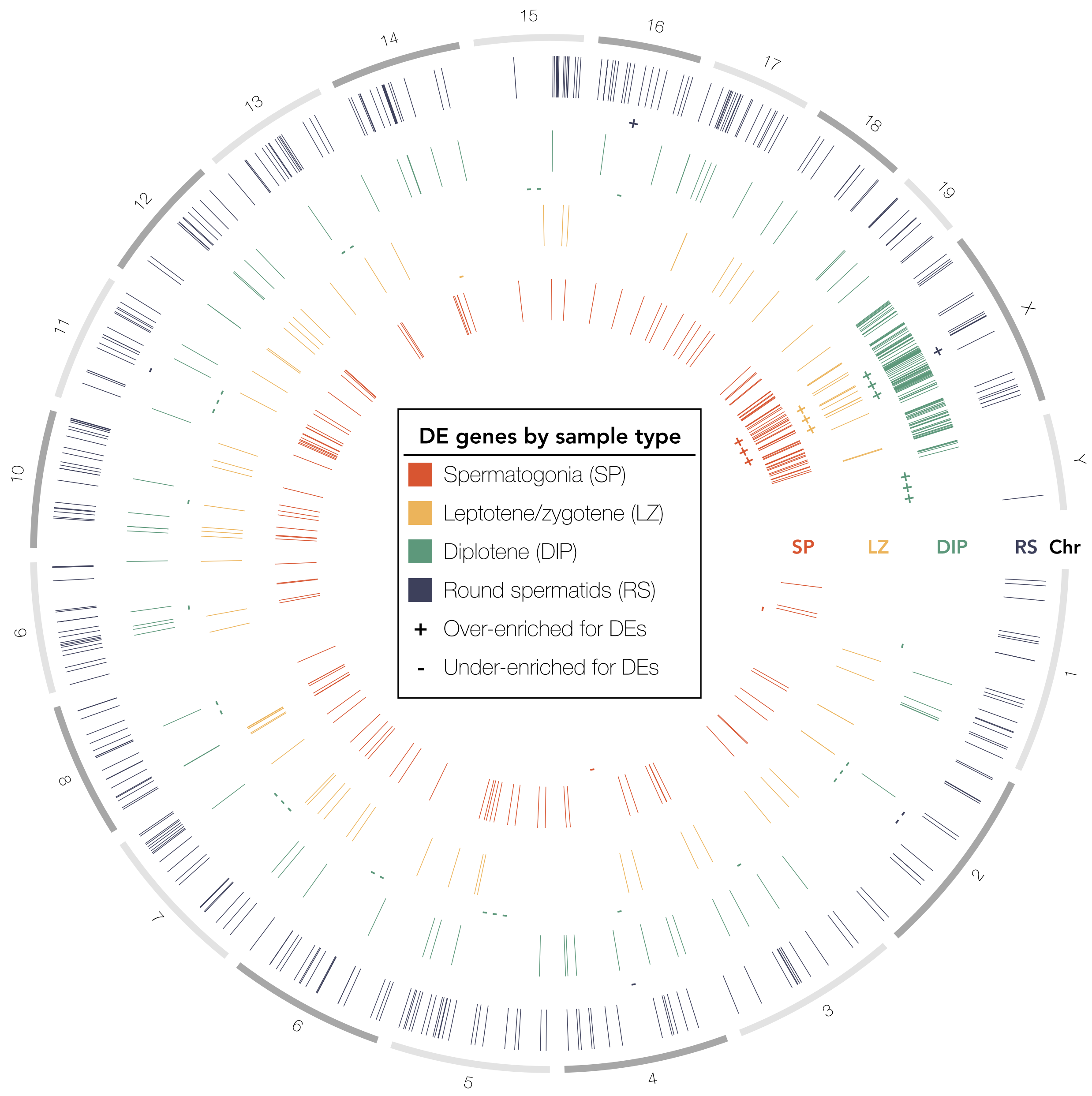


**Figure S9. Differential expression is enriched on the X chromosome in early and late spermatogenesis in house mouse hybrids.** Spatial distribution of transgressive DE genes across the reference mouse genome chromosomes (build GRCm38.p6) between sterile hybrid house mice and both parent crosses for spermatogonia (red), leptotene/zygotene spermatocytes (yellow), diplotene spermatocytes (green), and round spermatids (blue). Plus or minus symbols indicate whether a given chromosome has significant enrichment or depletion for DE genes. Number of plus or minus symbols indicates significance level according to hypergeometric tests (+/- indicates $p < 0.05$, ++/– indicates $p < 0.01$, +++/–- indicates $p < 0.001$).

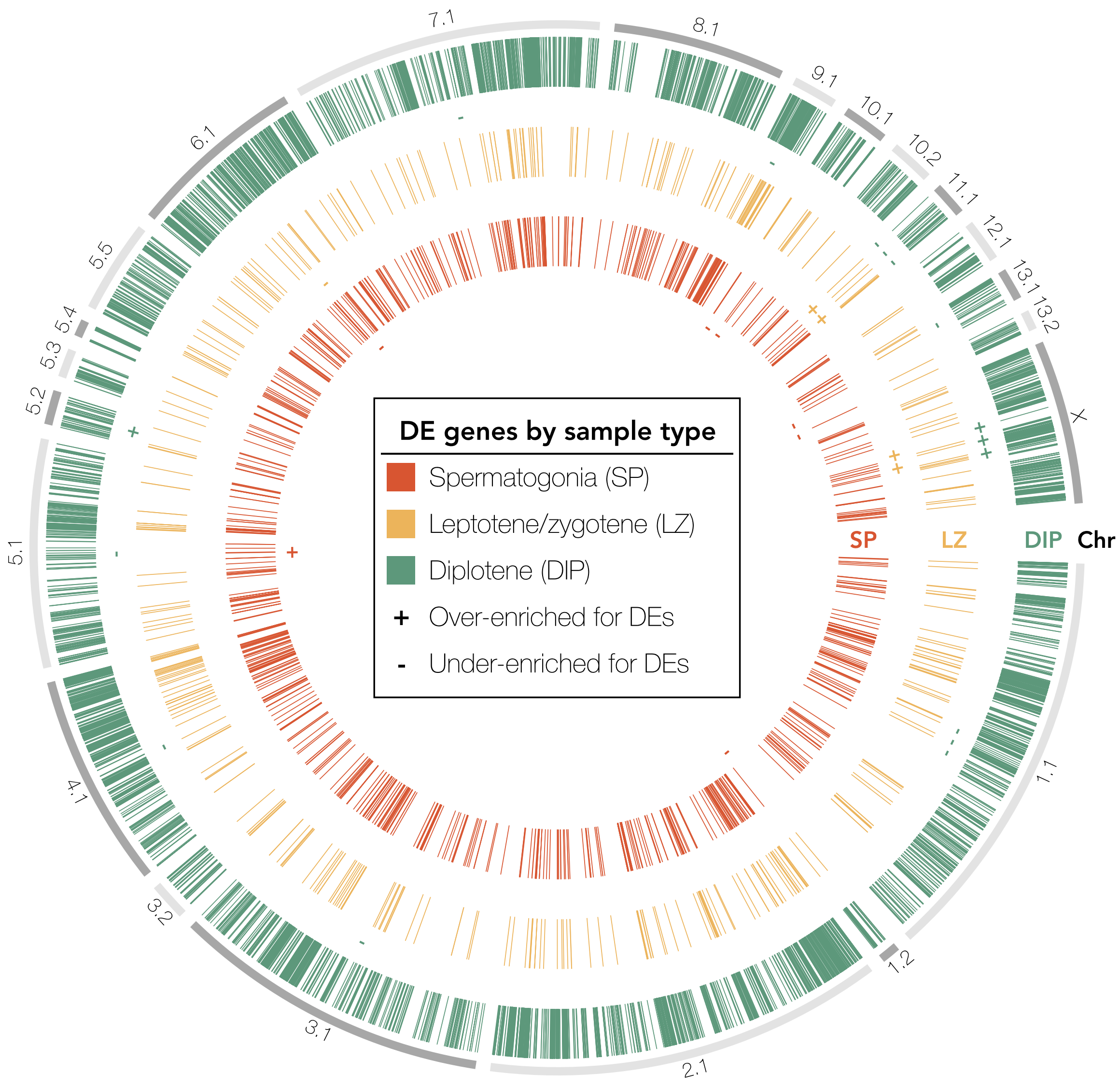


**Figure S10. Differential expression is only enriched on the X chromosome in later spermatogenesis in dwarf hamster hybrids.** Spatial distribution of transgressive DE genes across *P. sungorus* reference genome between hybrid dwarf hamsters and both parent species for spermatogonia (red), leptotene/zygotene spermatocytes (yellow), and diplotene spermatocytes (green). Plus or minus symbols indicate whether a given chromosome has significant enrichment or depletion for DE genes. Number of plus or minus symbols indicates significance level according to hypergeometric tests (+/- indicates p < 0.05, ++/– indicates p < 0.01, +++/–- indicates p < 0.001).

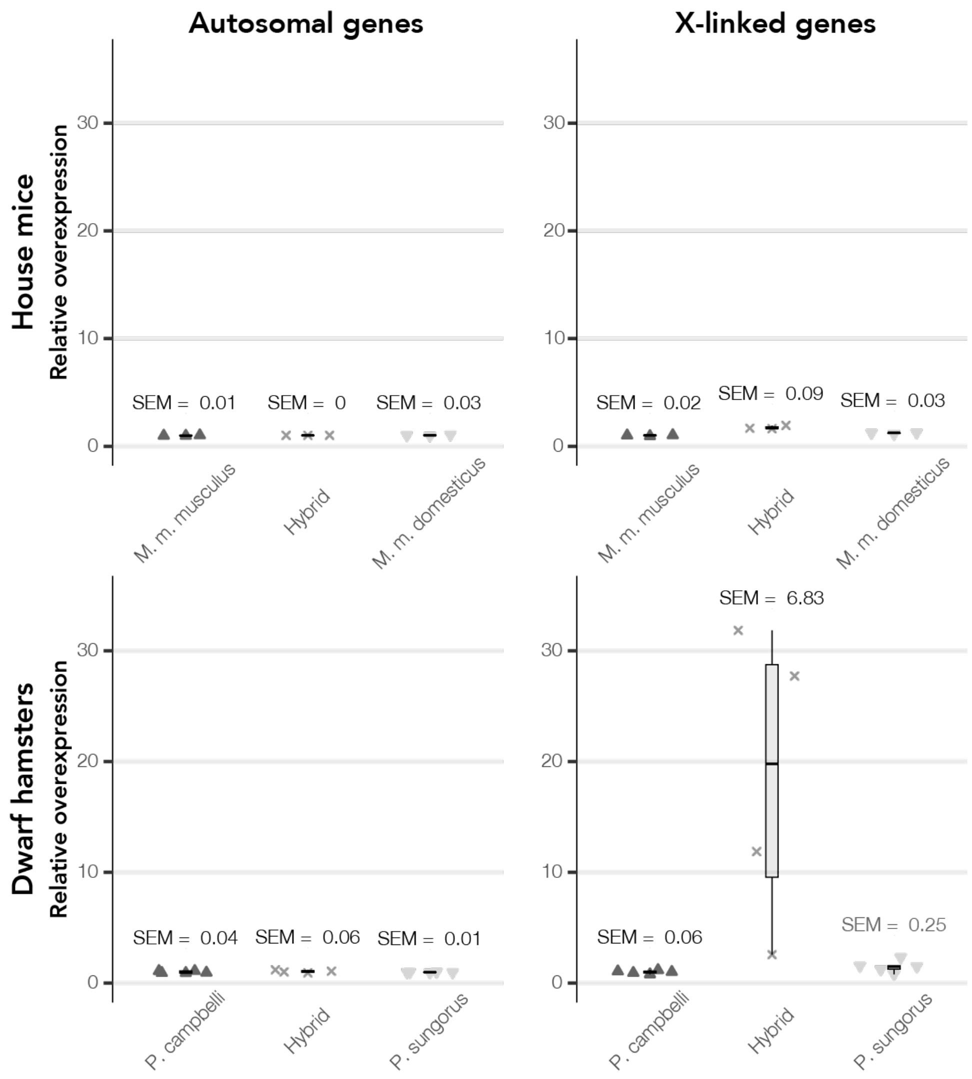


**Figure S11. Hybrid dwarf hamsters have extensive variability in the relative overexpression of X-linked genes during diplotene.** We assessed the extent of relative overexpression of autosomal (left panels) and X-linked genes (right panels) in hybrids compared to the mean expression (normalized RPKM) of X-linked genes during diplotene in parental males (*P. campbelli* for dwarf hamster and *M. m. musculus* for house mice). The standard error of the mean (SEM) is displayed above each box and whisker plot, and whiskers extend to either the largest or smallest value or no further than 1.5 times the interquartile range.

**Supplemental Tables:**

Table S1. Male reproductive traits of *P. campbelli*, *P. sungorus* and hybrids including body weight, paired testes weight, relative paired testes weight, seminal vesicle (SV) weight, relative SV weight, proportion motile sperm, and normalized sperm count.

| Species | Family | Individual | Age | Body weight (g) | Testes weight (mg) | Relative testes weight (mg/g) | SV weight (mg) | Relative SV Weight (mg/g) | Proportion motile sperm | Normalized sperm count (1 x 10^6) |
|---|---|---|---|---|---|---|---|---|---|---|
| BBBB | 153 | 5M | 103 | 34 | 1.76 | 5.17 | NA | NA | NA | NA |
| BBBB | 153 | 6M | 106 | 29.8 | 1.42 | 4.77 | 0.4 | 1.34 | NA | NA |
| BBBB | 154 | 4M | 111 | 30.1 | 1.39 | 4.62 | 0.43 | 1.43 | NA | NA |
| BBBB | 195 | 4M | 128 | 34.1 | 0.82 | 2.42 | 0.56 | NA | 0.79 | 1011 |
| BBBB | 197 | 3M | 141 | 36.2 | 1.47 | 4.04 | 0.54 | 1.49 | 0.92 | 675 |
| BBBB | 197 | 4M | 133 | 32 | 1.16 | 3.61 | 0.38 | 1.19 | 0.88 | 260 |
| BBBB | 198 | 7M | 104 | 35.9 | 1.59 | 4.44 | NA | NA | NA | NA |
| BBBB | 280 | 4M | 154 | 51.8 | 1.27 | 2.45 | 0.81 | 1.56 | NA | 1160 |
| BBBB | 281C | 3M | 83 | 30.5 | 1.05 | 3.43 | 0.38 | 1.26 | NA | 2240 |
| BBBB | 283B | 4M | 78 | 39.1 | 1.3 | 3.33 | 0.82 | 2.1 | NA | 1515 |
| BBBB | 291 | 3M | 163 | 43.9 | 1.39 | 3.16 | 0.6 | 1.36 | NA | NA |
| BBBB | 295A | 3M | 142 | 43.7 | 1.51 | 3.45 | 0.37 | NA | NA | NA |
| BBBB | 295A | 4M | NA | 41.8 | 1.35 | 3.23 | 0.47 | 1.12 | NA | NA |
| BBSS | 100 | 2M | 105 | 24.4 | 0.19 | 0.76 | 0.09 | 0.37 | 0 | 1 |
| BBSS | 7 | 1M | 85 | 26.8 | 0.24 | 0.9 | 0.09 | 0.34 | NA | 0 |
| BBSS | 90 | 2M | 97 | 20.8 | 0.19 | 0.9 | 0.07 | 0.34 | NA | 0 |
| BBSS | 90 | 3M | 105 | 14.1 | 0.04 | 0.31 | NA | NA | NA | 0 |
| BBSS | 95 | 3M | 81 | 18.3 | 0.05 | 0.25 | NA | NA | NA | 0 |
| BBSS | 99 | 3M | 116 | 20 | 0.13 | 0.64 | 0.09 | 0.47 | NA | 0 |
| SSSS | 138 | 5M | 200 | 32.7 | 0.66 | 2 | NA | NA | 0.65 | 217 |
| SSSS | 152 | 4M | 90 | 38.2 | 0.54 | 1.42 | 0.22 | 0.57 | NA | 335 |
| SSSS | 155 | 5M | 101 | 30.4 | 0.35 | 1.15 | 0.1 | 0.32 | 0.78 | 37 |
| SSSS | 155 | 6M | 73 | 31.7 | 0.43 | 1.34 | NA | NA | 0.78 | 142 |
| SSSS | 182 | 5M | 143 | 33.2 | 0.57 | 1.72 | 0.22 | NA | 0.91 | 1575 |
| SSSS | 188 | 4M | 127 | NA | 1.23 | NA | 0.34 | NA | 0.81 | 378 |
| SSSS | 188 | 5M | 140 | 39.9 | 0.81 | 2.02 | 0.32 | 0.81 | 0.78 | 250 |
| SSSS | 230 | 5M | 148 | 44.9 | 0.31 | 0.68 | 0.22 | 0.49 | NA | 1 |
| SSSS | 231 | 5M | 131 | 45.9 | 0.37 | 0.81 | 0.24 | 0.52 | NA | 0 |
| SSSS | 231 | 6M | 124 | 37.5 | 0.41 | 1.1 | 0.35 | 0.93 | NA | 59 |
| SSSS | 231 | 7M | 124 | 39.7 | 0.32 | 0.81 | 0.24 | 0.61 | NA | 0 |
| SSSS | 233A | 7M | 59 | 32 | 0.52 | 1.62 | 0.17 | 0.54 | NA | 220 |
| SSSS | 233B | 4M | 78 | 38.6 | 0.57 | 1.48 | 0.2 | 0.53 | NA | 370 |

Table S2. Sample information and read counts. Sample IDs correspond to the parent species or hybrid status of the individual (BBBB = P. campbelli, SSSS = P. sungorus, BBSS = hybrid), the individual ID number, and the sample type (SP = spermatogonia, LZ = leptotene/zygotene, DIP = diplotene, and RS = round spermatids).

| Species | Family | Individual | Cell Population | SRA Accession | Raw read pairs | Pcam-Alignment Reads (F+R) | Psun-Alignment Reads (F+R) | Post-Suspenders Reads (F+R) | Assigned featureCount reads (F+R) |
|---|---|---|---|---|---|---|---|---|---|
| BBBB | 153 | 5M | DIP | SRR26402739 | 28704759 | 36718525 | 48604390 | 49096504 | 36144982 |
| BBBB | 153 | 5M | LZ | SRR26402738 | 29761942 | 52272791 | 51392219 | 52268383 | 39635287 |
| BBBB | 153 | 5M | SP | SRR26402727 | 32343248 | 57045203 | 55846437 | 56967393 | 41891105 |
| BBBB | 153 | 6M | DIP | SRR26402716 | 29117549 | 49411834 | 48652305 | 49381241 | 36817154 |
| BBBB | 153 | 6M | SP | SRR26402705 | 25010611 | 41242411 | 40470003 | 41173619 | 31032766 |
| BBBB | 154 | 4M | LZ | SRR26402698 | 27439376 | 47729051 | 46779555 | 47679328 | 35053421 |
| BBBB | 195 | 4M | DIP | SRR26402697 | 28654583 | 49990063 | 49120565 | 49938023 | 36328447 |
| BBBB | 195 | 4M | LZ | SRR26402696 | 24757971 | 41326550 | 40607666 | 41297838 | 30787675 |
| BBBB | 195 | 4M | RS | SRR26402695 | 33949981 | 59348339 | 58448456 | 59266886 | 44663398 |
| BBBB | 195 | 4M | SP | SRR26402694 | 32744540 | 57038191 | 55911864 | 57004331 | 42052840 |
| BBBB | 197 | 3M | DIP | SRR26402737 | 28774386 | 48107812 | 47353483 | 48093420 | 35753106 |
| BBBB | 197 | 3M | RS | SRR26402736 | 27449504 | 46265757 | 45565843 | 46282313 | 34011365 |
| BBBB | 197 | 4M | DIP | SRR26402735 | 20293025 | 35136632 | 34580893 | 35103192 | 26313579 |
| BBBB | 197 | 4M | LZ | SRR26402734 | 27698727 | 48025107 | 47162803 | 47986743 | 35773804 |
| BBBB | 197 | 4M | RS | SRR26402733 | 35077355 | 61171415 | 60233939 | 61071343 | 45923141 |
| BBBB | 198 | 7M | LZ | SRR26402732 | 34852328 | 60621717 | 59584169 | 60529076 | 46006840 |
| BBBB | 198 | 7M | RS | SRR26402731 | 34654461 | 59578475 | 58568459 | 59515442 | 44093846 |
| BBBB | 198 | 7M | SP | SRR26402730 | 20593687 | 36069646 | 35391134 | 36061839 | 26589436 |
| BBSS | 100 | 2M | DIP | SRR26402729 | 27602009 | 47830055 | 47867756 | 48102298 | 34916044 |
| BBSS | 100 | 2M | LZ | SRR26402728 | 24877481 | 43944813 | 43945348 | 44194298 | 33154338 |
| BBSS | 100 | 2M | SP | SRR26402726 | 28894633 | 50089417 | 50074357 | 50326541 | 38236146 |
| BBSS | 90 | 2M | DIP | SRR26402725 | 23483730 | 40776752 | 40816383 | 41048955 | 29872793 |
| BBSS | 90 | 2M | LZ | SRR26402724 | 29785440 | 52471629 | 52517663 | 52853160 | 38545691 |
| BBSS | 90 | 2M | SP | SRR26402723 | 21232899 | 36971419 | 36954811 | 37215967 | 26873790 |
| BBSS | 90 | 3M | SP | SRR26402722 | 26505885 | 46063910 | 46056188 | 46381121 | 33794829 |

| | | | | | | | | | |
|---|---|---|---|---|---|---|---|---|---|
| BBSS | 95 | 3M | DIP | SRR26402721 | 25731045 | 45036605 | 45099780 | 45371762 | 32527867 |
| BBSS | 95 | 3M | LZ | SRR26402720 | 23113566 | 40667672 | 40691223 | 40942063 | 29784990 |
| BBSS | 99 | 3M | DIP | SRR26402719 | 32412672 | 57034662 | 57048691 | 57417683 | 42164444 |
| BBSS | 99 | 3M | LZ | SRR26402718 | 29218077 | 49191580 | 49179410 | 49444223 | 37409477 |
| BBSS | 99 | 3M | SP | SRR26402717 | 26752102 | 46674332 | 46642011 | 46909673 | 35243220 |
| SSSS | 138 | 5M | DIP | SRR26402715 | 31668197 | 55539097 | 56518165 | 56130476 | 41754867 |
| SSSS | 138 | 5M | LZ | SRR26402714 | 30150190 | 53563250 | 54626186 | 54330554 | 38860722 |
| SSSS | 155 | 5M | DIP | SRR26402713 | 27927121 | 49231857 | 50096727 | 49771221 | 36336601 |
| SSSS | 155 | 5M | LZ | SRR26402712 | 31211602 | 54221237 | 55341593 | 55014305 | 39108953 |
| SSSS | 155 | 5M | SP | SRR26402711 | 31212480 | 54501098 | 55527020 | 55174124 | 41077419 |
| SSSS | 155 | 6M | DIP | SRR26402710 | 34881395 | 62036149 | 63214304 | 62744812 | 46111924 |
| SSSS | 155 | 6M | LZ | SRR26402709 | 28282870 | 48212032 | 49274841 | 48978992 | 35117846 |
| SSSS | 155 | 6M | SP | SRR26402708 | 29366707 | 50059914 | 50979161 | 50701265 | 37591448 |
| SSSS | 182 | 5M | DIP | SRR26402707 | 29552280 | 52234804 | 53192430 | 52832743 | 39037762 |
| SSSS | 182 | 5M | LZ | SRR26402706 | 28835525 | 49447409 | 50441610 | 50168805 | 34229672 |
| SSSS | 182 | 5M | RS | SRR26402704 | 29208957 | 49251459 | 50247066 | 49947638 | 35996988 |
| SSSS | 188 | 4M | DIP | SRR26402703 | 38506160 | 67405285 | 68807300 | 68349156 | 48314454 |
| SSSS | 188 | 4M | RS | SRR26402702 | 23229053 | 40549254 | 41398361 | 41122282 | 29867971 |
| SSSS | 188 | 4M | SP | SRR26402701 | 29676568 | 52766550 | 53818028 | 53503297 | 38871363 |
| SSSS | 188 | 5M | RS | SRR26402700 | 24605003 | 43499551 | 44267170 | 43967979 | 33273278 |
| SSSS | 188 | 5M | SP | SRR26402699 | 21803869 | 38100344 | 38926447 | 38682327 | 28078102 |

Table S3. Summary of differential expression (DE) of genes in the PAR between hybrids and P. campbelli and hybrids and P. sungorus during spermatogonia (SP), leptotene/zygotene (LZ), and diplotene (DIP). For each gene, the homologous M. musculus (GRCm38.p6) gene, the start position, the stop position, and whether the gene was DE between hybrids and parents in a given stage are given.

| ***P. sungorus* gene** | **Mouse Ensembl ID** | **Start (bp)** | **Stop (bp)** | **GRCm38.p6 gene** | **DE between hybrids and *P. cam* in SP?** | **DE between hybrids and *P. sun* in SP?** | **DE between hybrids and *P. cam* in LZ?** | **DE between hybrids and *P. sun* in LZ?** | **DE between hybrids and *P. cam* in DIP?** | **DE between hybrids and *P. sun* in DIP?** |
|---|---|---|---|---|---|---|---|---|---|---|
| Psun_G000022875 | ENSMUSG00000025607 | 115715330 | 115715572 | NA | | | | | | |
| Psun_G000022876 | ENSMUSG00000008813 | 115795915 | 115796010 | Tppp2 | | | | | | |
| Psun_G000022877 | ENSMUSG00000068392 | 115947535 | 115947780 | NA | | | | | | |
| Psun_G000022878 | ENSMUSG00000069227 | 115949671 | 115949842 | Gprin1 | X | X | X | | | |
| Psun_G000022879 | ENSMUSG00000004558 | 115957028 | 115957314 | Ndrg2 | | X | | | X | X |
| Psun_G000022880 | NA | 116147271 | 116147334 | NA | | | | | | |
| Psun_G000022881 | ENSMUSG00000029088 | 116595188 | 116595208 | Kcnip4 | | | | | | |
| Psun_G000022882 | ENSMUSG00000004558 | 116752226 | 116752329 | Ndrg2 | | | | | | |
| Psun_G000022883 | ENSMUSG00000053293 | 117437385 | 117438166 | NA | | | | | X | |
| Psun_G000022884 | NA | 117846192 | 117846210 | NA | | | | | | |
| Psun_G000022885 | ENSMUSG00000053293 | 117846621 | 117846746 | NA | | | | | | |
| Psun_G000022886 | ENSMUSG00000053293 | 118328521 | 118328813 | NA | | | | | | |
| Psun_G000022887 | ENSMUSG00000053293 | 118331294 | 118331305 | NA | | | | | | |
| Psun_G000022888 | ENSMUSG00000053347 | 118332594 | 118332612 | NA | | | | | | |
| Psun_G000022889 | ENSMUSG00000053465 | 118341279 | 118341470 | Hs6st3 | | | | | | |

Table S4. Results from tests determining whether there are more (Enrichment P-Value) or fewer (Depletion P-value) shared transgressive DE genes between hybrid house mice and hybrid dwarf hamsters for each stage of spermatogenesis than expected by chance. Enrichment or depletion was determined using the *phyper* function in R, and the expected versus observed amount of overlap, as well as the number of house mouse hybrid or dwarf hamster hybrid genes alone are included.

| Stage | Depletion P-Value | Enrichment P-Value | Expected Overlap | Observed Overlap | Number of Mouse DE Genes | Number of Hamster DE Genes |
|---|---|---|---|---|---|---|
| Spermatogonia | 0.40 | 0.82 | 3 | 2 | 53 | 662 |
| Leptotene/zygotene | 0.88 | 0.45 | 1 | 1 | 21 | 322 |
| Diplotene | 1.00 | > 0.001 | 53 | 68 | 103 | 3830 |

Table S5. House mouse Ensembl IDs, gene descriptions, and gene symbols of shared transgressive DE genes between hybrid house mice and hybrid dwarf hamsters for each stage of spermatogenesis.

| Mouse Ensembl ID | Gene Description | Gene Symbol | Stage |
|---|---|---|---|
| ENSMUSG00000047281 | stratifin | Sfn | Spermatogonia |
| ENSMUSG00000062168 | protein phosphatase with EF hand calcium-binding domain 1 | Ppef1 | Spermatogonia |
| ENSMUSG00000035187 | NK6 homeobox 1 | Nkx6-1 | Leptotene/zygotene |
| ENSMUSG00000038855 | inositol 1,4,5-trisphosphate 3-kinase B | Itpkb | Diplotene |
| ENSMUSG00000002015 | B cell receptor associated protein 31 | Bcap31 | Diplotene |
| ENSMUSG00000079642 | cancer/testis antigen 47 | Ct47 | Diplotene |
| ENSMUSG00000073207 | coiled-coil domain containing 160 | Ccdc160 | Diplotene |
| ENSMUSG00000082728 | SPT20 SAGA complex component, pseudogene | Supt20-ps | Diplotene |
| ENSMUSG00000031347 | centrin 2 | Cetn2 | Diplotene |
| ENSMUSG00000043453 | MAGE family member A10 | Magea10 | Diplotene |
| ENSMUSG00000031432 | phosphoribosyl pyrophosphate synthetase 1 | Prps1 | Diplotene |
| ENSMUSG00000025289 | peroxiredoxin 4 | Prdx4 | Diplotene |
| ENSMUSG00000067649 | MAGE family member B18 | Mageb18 | Diplotene |
| ENSMUSG00000009596 | TATA-box binding protein associated factor 7 like | Taf7l | Diplotene |
| ENSMUSG00000096966 | PNMA family member 6E | Pnma6e | Diplotene |
| ENSMUSG00000009941 | nuclear RNA export factor 2 | Nxf2 | Diplotene |
| ENSMUSG00000031176 | dynein light chain Tctex-type 3 | Dynlt3 | Diplotene |
| ENSMUSG00000031007 | ATPase, H+ transporting, lysosomal accessory protein 2 | Atp6ap2 | Diplotene |
| ENSMUSG00000031198 | FUN14 domain containing 2 | Fundc2 | Diplotene |
| ENSMUSG00000031155 | proviral integration site 2 | Pim2 | Diplotene |
| ENSMUSG00000026641 | upstream transcription factor 1 | Usf1 | Diplotene |
| ENSMUSG00000021384 | sushi domain containing 3 | Susd3 | Diplotene |
| ENSMUSG00000031349 | NAD(P) dependent steroid dehydrogenase-like | Nsdhl | Diplotene |
| ENSMUSG00000031358 | MSL complex subunit 3 | Msl3 | Diplotene |
| ENSMUSG00000067878 | MAP7 domain containing 3 | Map7d3 | Diplotene |
| ENSMUSG00000043549 | family with sequence similarity 90, member A1B | Fam90a1b | Diplotene |
| ENSMUSG00000002014 | signal sequence receptor, delta | Ssr4 | Diplotene |
| ENSMUSG00000042271 | nuclear transport factor 2-like export factor 2 | Nxt2 | Diplotene |
| ENSMUSG00000025268 | MAGE family member D2 | Maged2 | Diplotene |
| ENSMUSG00000067430 | zinc finger protein 763 | Zfp763 | Diplotene |
| ENSMUSG00000002274 | meteorin, glial cell differentiation regulator | Metrn | Diplotene |
| ENSMUSG00000025264 | TSR2 20S rRNA accumulation | Tsr2 | Diplotene |
| ENSMUSG00000031095 | cullin 4B | Cul4b | Diplotene |
| ENSMUSG00000032750 | growth factor receptor bound protein 2-associated protein 3 | Gab3 | Diplotene |
| ENSMUSG00000071719 | NALCN channel auxiliary factor 2 | Nalf2 | Diplotene |
| ENSMUSG00000041020 | MAP7 domain containing 2 | Map7d2 | Diplotene |
| ENSMUSG00000031403 | dyskeratosis congenita 1, dyskerin | Dkc1 | Diplotene |
| ENSMUSG00000025037 | monoamine oxidase A | Maoa | Diplotene |
| ENSMUSG00000031134 | RNA binding motif protein, X chromosome | Rbmx | Diplotene |
| ENSMUSG00000031431 | TSC22 domain family, member 3 | Tsc22d3 | Diplotene |
| ENSMUSG00000045427 | heterogeneous nuclear ribonucleoprotein H2 | Hnrnph2 | Diplotene |

| | | | |
|---|---|---|---|
| ENSMUSG00000041064 | PIF1 5'-to-3' DNA helicase | Pif1 | Diplotene |
| ENSMUSG00000036022 | PABIR family member 2 | Pabir2 | Diplotene |
| ENSMUSG00000081133 | reproductive homeobox 11, pseudogene 2 | Rhox11-ps2 | Diplotene |
| ENSMUSG00000051592 | cyclin B3 | Ccnb3 | Diplotene |
| ENSMUSG00000031066 | ubiquitin specific peptidase 11 | Usp11 | Diplotene |
| ENSMUSG00000072964 | basic helix-loop-helix domain containing, class B9 | Bhlhb9 | Diplotene |
| ENSMUSG00000067873 | HIV TAT specific factor 1 | Htatsf1 | Diplotene |
| ENSMUSG00000031367 | adaptor-related protein complex 1, sigma 2 subunit | Ap1s2 | Diplotene |
| ENSMUSG00000037315 | jade family PHD finger 3 | Jade3 | Diplotene |
| ENSMUSG00000051220 | excision repair cross-complementing rodent repair deficiency complementation group 6 like | Ercc6l | Diplotene |
| ENSMUSG00000031805 | Janus kinase 3 | Jak3 | Diplotene |
| ENSMUSG00000034311 | kinesin family member 4 | Kif4 | Diplotene |
| ENSMUSG00000015214 | myotubularin related protein 1 | Mtmr1 | Diplotene |
| ENSMUSG00000000037 | Scm polycomb group protein like 2 | Scml2 | Diplotene |
| ENSMUSG00000034055 | phosphorylase kinase alpha 1 | Phka1 | Diplotene |
| ENSMUSG00000033737 | fibronectin type III domain containing 3C1 | Fndc3c1 | Diplotene |
| ENSMUSG00000001127 | Araf proto-oncogene, serine/threonine kinase | Araf | Diplotene |
| ENSMUSG00000025246 | transducin (beta)-like 1 X-linked | Tbl1x | Diplotene |
| ENSMUSG00000047694 | Yip1 domain family, member 6 | Yipf6 | Diplotene |
| ENSMUSG00000025272 | trophinin | Tro | Diplotene |
| ENSMUSG00000062949 | ATPase, class VI, type 11C | Atp11c | Diplotene |
| ENSMUSG00000079487 | mediator complex subunit 12 | Med12 | Diplotene |
| ENSMUSG00000079509 | zinc finger protein X-linked | Zfx | Diplotene |
| ENSMUSG00000079532 | CTAG2 like 2 | Ctag2l2 | Diplotene |
| ENSMUSG00000031310 | zinc finger, MYM-type 3 | Zmym3 | Diplotene |
| ENSMUSG00000057421 | LAS1-like (S. cerevisiae) | Las1l | Diplotene |
| ENSMUSG00000036109 | muscleblind like splicing factor 3 | Mbnl3 | Diplotene |
| ENSMUSG00000056537 | ring finger protein, LIM domain interacting | Rlim | Diplotene |
| ENSMUSG00000000838 | fragile X messenger ribonucleoprotein 1 | Fmr1 | Diplotene |
| ENSMUSG00000031059 | NADH:ubiquinone oxidoreductase subunit B11 | Ndufb11 | Diplotene |
| Mouse Ensembl ID | Gene Description | Gene Symbol | Stage |
| ENSMUSG00000047281 | stratifin | Sfn | Spermatogonia |
| ENSMUSG00000062168 | protein phosphatase with EF hand calcium-binding domain 1 | Ppef1 | Spermatogonia |
| ENSMUSG00000035187 | NK6 homeobox 1 | Nkx6-1 | Leptotene/zygotene |
| ENSMUSG00000038855 | inositol 1,4,5-trisphosphate 3-kinase B | Itpkb | Diplotene |
| ENSMUSG00000002015 | B cell receptor associated protein 31 | Bcap31 | Diplotene |
| ENSMUSG00000079642 | cancer/testis antigen 47 | Ct47 | Diplotene |
| ENSMUSG00000073207 | coiled-coil domain containing 160 | Ccdc160 | Diplotene |
| ENSMUSG00000082728 | SPT20 SAGA complex component, pseudogene | Supt20-ps | Diplotene |
| ENSMUSG00000031347 | centrin 2 | Cetn2 | Diplotene |
| ENSMUSG00000043453 | MAGE family member A10 | Magea10 | Diplotene |

Table S6. Enrichment P-values, term sizes, number of DE genes included in tests, intersection size between term sizes and number of DE genes, gene background size, and GO Term information including Term ID, Source, and Name are shown for GO Terms with significant enrichment amongst transgressive DE genes in hybrid dwarf hamster spermatogonia samples. Gene backgrounds were the number of genes “expressed” in hybrids and both parent species. Enrichment was determined using the gProfiler2 package in R (Kolberg *et al.* 2020).

| P-Value | Term Size | Query Size (Number DE) | Intersection Size | Background Size | GO Term ID | GO Term Source | GO Term Name |
|---|---|---|---|---|---|---|---|
| 7.25E-41 | 178 | 755 | 73 | 13823 | GO:0003341 | GO:BP | cilium movement |
| 5.81E-35 | 147 | 755 | 62 | 13823 | GO:0060285 | GO:BP | cilium-dependent cell motility |
| 5.81E-35 | 147 | 755 | 62 | 13823 | GO:0001539 | GO:BP | cilium or flagellum-dependent cell motility |
| 1.10E-34 | 143 | 755 | 61 | 13823 | GO:0060294 | GO:BP | cilium movement involved in cell motility |
| 1.10E-29 | 124 | 755 | 53 | 13823 | GO:0030317 | GO:BP | flagellated sperm motility |
| 1.10E-29 | 124 | 755 | 53 | 13823 | GO:0097722 | GO:BP | sperm motility |
| 7.94E-25 | 502 | 755 | 98 | 13823 | GO:0048232 | GO:BP | male gamete generation |
| 1.20E-24 | 486 | 755 | 96 | 13823 | GO:0007283 | GO:BP | spermatogenesis |
| 2.52E-24 | 358 | 755 | 81 | 13823 | GO:0007018 | GO:BP | microtubule-based movement |
| 2.18E-20 | 801 | 755 | 120 | 13823 | GO:0019953 | GO:BP | sexual reproduction |
| 4.13E-19 | 622 | 755 | 101 | 13823 | GO:0007276 | GO:BP | gamete generation |
| 1.31E-18 | 180 | 755 | 51 | 13823 | GO:0048515 | GO:BP | spermatid differentiation |
| 1.78E-18 | 174 | 755 | 50 | 13823 | GO:0007286 | GO:BP | spermatid development |
| 2.17E-18 | 754 | 755 | 112 | 13823 | GO:0032504 | GO:BP | multicellular organism reproduction |
| 3.23E-18 | 88 | 755 | 36 | 13823 | GO:0035082 | GO:BP | axoneme assembly |
| 1.44E-17 | 716 | 755 | 107 | 13823 | GO:0048609 | GO:BP | multicellular organismal reproductive process |
| 3.36E-16 | 354 | 755 | 69 | 13823 | GO:0044782 | GO:BP | cilium organization |
| 4.66E-16 | 118 | 755 | 39 | 13823 | GO:0001578 | GO:BP | microtubule bundle formation |
| 6.60E-16 | 1132 | 755 | 139 | 13823 | GO:0022414 | GO:BP | reproductive process |
| 1.17E-15 | 1139 | 755 | 139 | 13823 | GO:0000003 | GO:BP | reproduction |
| 4.72E-14 | 329 | 755 | 63 | 13823 | GO:0060271 | GO:BP | cilium assembly |
| 2.03E-12 | 777 | 755 | 102 | 13823 | GO:0003006 | GO:BP | developmental process involved in reproduction |
| 5.88E-12 | 391 | 755 | 66 | 13823 | GO:0022412 | GO:BP | cellular process involved in reproduction in multicellular organism |
| 6.44E-12 | 790 | 755 | 102 | 13823 | GO:0007017 | GO:BP | microtubule-based process |
| 4.49E-11 | 298 | 755 | 55 | 13823 | GO:0007281 | GO:BP | germ cell development |
| 2.58E-09 | 62 | 755 | 23 | 13823 | GO:0044458 | GO:BP | motile cilium assembly |
| 3.11E-07 | 500 | 755 | 67 | 13823 | GO:0030031 | GO:BP | cell projection assembly |
| 3.92E-07 | 491 | 755 | 66 | 13823 | GO:0120031 | GO:BP | plasma membrane bounded cell projection assembly |
| 7.65E-06 | 41 | 755 | 16 | 13823 | GO:0120316 | GO:BP | sperm flagellum assembly |
| 1.17E-05 | 42 | 755 | 16 | 13823 | GO:0003351 | GO:BP | epithelial cilium movement involved in extracellular fluid movement |

| | | | | | | | |
|---|---|---|---|---|---|---|---|
| 3.90E-05 | 45 | 755 | 16 | 13823 | GO:0006858 | GO:BP | extracellular transport |
| 0.00011694 | 124 | 755 | 26 | 13823 | GO:0009566 | GO:BP | fertilization |
| 0.00130436 | 86 | 755 | 20 | 13823 | GO:0007338 | GO:BP | single fertilization |
| 0.00218032 | 26 | 755 | 11 | 13823 | GO:0007288 | GO:BP | sperm axoneme assembly |
| 0.0058181 | 34 | 755 | 12 | 13823 | GO:0070286 | GO:BP | axonemal dynein complex assembly |
| 0.00732699 | 10 | 755 | 7 | 13823 | GO:0007618 | GO:BP | mating |
| 2.72E-47 | 241 | 755 | 90 | 13823 | GO:0031514 | GO:CC | motile cilium |
| 1.05E-42 | 180 | 755 | 75 | 13823 | GO:0097729 | GO:CC | 9+2 motile cilium |
| 3.72E-37 | 158 | 755 | 66 | 13823 | GO:0036126 | GO:CC | sperm flagellum |
| 9.52E-34 | 613 | 755 | 123 | 13823 | GO:0005929 | GO:CC | cilium |
| 9.00E-24 | 156 | 755 | 53 | 13823 | GO:0005930 | GO:CC | axoneme |
| 9.00E-24 | 156 | 755 | 53 | 13823 | GO:0097014 | GO:CC | ciliary plasm |
| 1.68E-14 | 129 | 755 | 39 | 13823 | GO:0001669 | GO:CC | acrosomal vesicle |
| 3.31E-14 | 247 | 755 | 54 | 13823 | GO:0032838 | GO:CC | plasma membrane bounded cell projection cytoplasm |
| 7.06E-12 | 277 | 755 | 54 | 13823 | GO:0099568 | GO:CC | cytoplasmic region |
| 1.46E-10 | 36 | 755 | 19 | 13823 | GO:0005879 | GO:CC | axonemal microtubule |
| 1.40E-08 | 34 | 755 | 17 | 13823 | GO:0097228 | GO:CC | sperm principal piece |
| 1.39E-06 | 1965 | 755 | 173 | 13823 | GO:0042995 | GO:CC | cell projection |
| 2.95E-06 | 1893 | 755 | 167 | 13823 | GO:0120025 | GO:CC | plasma membrane bounded cell projection |
| 5.69E-05 | 46 | 755 | 16 | 13823 | GO:0097225 | GO:CC | sperm midpiece |
| 0.00012112 | 13 | 755 | 9 | 13823 | GO:0001534 | GO:CC | radial spoke |
| 0.00115749 | 303 | 755 | 42 | 13823 | GO:0030141 | GO:CC | secretory granule |
| 0.01260294 | 30 | 755 | 11 | 13823 | GO:0002080 | GO:CC | acrosomal membrane |
| 0.01791024 | 91 | 755 | 19 | 13823 | GO:0005881 | GO:CC | cytoplasmic microtubule |
| 0.04336422 | 47 | 755 | 13 | 13823 | GO:0030286 | GO:CC | dynein complex |
| 0.04389616 | 12 | 755 | 7 | 13823 | GO:0097224 | GO:CC | sperm connecting piece |
| 0.04389616 | 12 | 755 | 7 | 13823 | GO:0036128 | GO:CC | CatSper complex |

Table S7. Enrichment P-values, term sizes, number of DE genes included in tests, intersection size between term sizes and number of DE genes, gene background size, and GO Term information including Term ID, Source, and Name are shown for GO Terms with significant enrichment amongst transgressive DE genes in hybrid dwarf hamster leptotene/zygotene samples. Gene backgrounds were the number of genes "expressed" in hybrids and both parent species. Enrichment was determined using the gProfiler2 package in R (Kolberg *et al.* 2020).

| P-Value | Term Size | Query Size (Number DE) | Intersection Size | Background Size | GO Term ID | GO Term Source | GO Term Name |
|---|---|---|---|---|---|---|---|
| 9.51E-15 | 143 | 380 | 31 | 13788 | GO:0060294 | GO:BP | cilium movement involved in cell motility |
| 2.24E-14 | 147 | 380 | 31 | 13788 | GO:0060285 | GO:BP | cilium-dependent cell motility |
| 2.24E-14 | 147 | 380 | 31 | 13788 | GO:0001539 | GO:BP | cilium or flagellum-dependent cell motility |
| 1.02E-13 | 178 | 380 | 33 | 13788 | GO:0003341 | GO:BP | cilium movement |
| 1.68E-08 | 124 | 380 | 23 | 13788 | GO:0030317 | GO:BP | flagellated sperm motility |
| 1.68E-08 | 124 | 380 | 23 | 13788 | GO:0097722 | GO:BP | sperm motility |
| 1.44E-07 | 485 | 380 | 44 | 13788 | GO:0007283 | GO:BP | spermatogenesis |
| 4.31E-07 | 501 | 380 | 44 | 13788 | GO:0048232 | GO:BP | male gamete generation |
| 3.64E-06 | 358 | 380 | 35 | 13788 | GO:0007018 | GO:BP | microtubule-based movement |
| 1.34E-05 | 800 | 380 | 55 | 13788 | GO:0019953 | GO:BP | sexual reproduction |
| 0.000108749 | 753 | 380 | 51 | 13788 | GO:0032504 | GO:BP | multicellular organism reproduction |
| 0.000134217 | 621 | 380 | 45 | 13788 | GO:0007276 | GO:BP | gamete generation |
| 0.000175096 | 1131 | 380 | 66 | 13788 | GO:0022414 | GO:BP | reproductive process |
| 0.000224411 | 1138 | 380 | 66 | 13788 | GO:0000003 | GO:BP | reproduction |
| 0.001260874 | 715 | 380 | 47 | 13788 | GO:0048609 | GO:BP | multicellular organismal reproductive process |
| 0.03038005 | 86 | 380 | 13 | 13788 | GO:0007338 | GO:BP | single fertilization |
| 0.036247298 | 776 | 380 | 46 | 13788 | GO:0003006 | GO:BP | developmental process involved in reproduction |
| 1.51E-17 | 179 | 380 | 37 | 13788 | GO:0097729 | GO:CC | 9+2 motile cilium |
| 1.81E-17 | 240 | 380 | 42 | 13788 | GO:0031514 | GO:CC | motile cilium |
| 1.75E-14 | 157 | 380 | 32 | 13788 | GO:0036126 | GO:CC | sperm flagellum |
| 1.07E-09 | 611 | 380 | 54 | 13788 | GO:0005929 | GO:CC | cilium |
| 1.76E-05 | 129 | 380 | 20 | 13788 | GO:0001669 | GO:CC | acrosomal vesicle |
| 6.95E-05 | 35 | 380 | 11 | 13788 | GO:0005879 | GO:CC | axonemal microtubule |
| 8.16E-05 | 155 | 380 | 21 | 13788 | GO:0097014 | GO:CC | ciliary plasm |
| 8.16E-05 | 155 | 380 | 21 | 13788 | GO:0005930 | GO:CC | axoneme |

Table S8. Enrichment P-values, term sizes, number of DE genes included in tests, intersection size between term sizes and number of DE genes, gene background size, and GO Term information including Term ID, Source, and Name are shown for GO Terms with significant enrichment amongst transgressive DE genes in hybrid dwarf hamster diplotene samples. Gene backgrounds were the number of genes "expressed" in hybrids and both parent species. Enrichment was determined using the gProfiler2 package in R (Kolberg *et al.* 2020).

| P-Value | Term Size | Query Size (Number DE) | Intersection Size | Background Size | GO Term ID | GO Term Source | GO Term Name |
|---|---|---|---|---|---|---|---|
| 1.59E-33 | 2891 | 4583 | 1255 | 13797 | GO:0048731 | GO:BP | system development |
| 3.27E-26 | 3380 | 4583 | 1400 | 13797 | GO:0007275 | GO:BP | multicellular organism development |
| 3.50E-25 | 1987 | 4583 | 884 | 13797 | GO:0009653 | GO:BP | anatomical structure morphogenesis |
| 7.02E-25 | 4189 | 4583 | 1680 | 13797 | GO:0048856 | GO:BP | anatomical structure development |
| 9.96E-22 | 4945 | 4583 | 1925 | 13797 | GO:0032501 | GO:BP | multicellular organismal process |
| 1.40E-21 | 4542 | 4583 | 1785 | 13797 | GO:0032502 | GO:BP | developmental process |
| 2.17E-21 | 3171 | 4583 | 1301 | 13797 | GO:0030154 | GO:BP | cell differentiation |
| 2.55E-21 | 3172 | 4583 | 1301 | 13797 | GO:0048869 | GO:BP | cellular developmental process |
| 5.10E-19 | 2180 | 4583 | 929 | 13797 | GO:0048513 | GO:BP | animal organ development |
| 7.03E-19 | 883 | 4583 | 432 | 13797 | GO:0031175 | GO:BP | neuron projection development |
| 5.59E-18 | 1859 | 4583 | 805 | 13797 | GO:0007399 | GO:BP | nervous system development |
| 5.77E-18 | 953 | 4583 | 457 | 13797 | GO:0007155 | GO:BP | cell adhesion |
| 1.00E-17 | 1838 | 4583 | 796 | 13797 | GO:0007166 | GO:BP | cell surface receptor signaling pathway |
| 1.76E-17 | 1841 | 4583 | 796 | 13797 | GO:0050793 | GO:BP | regulation of developmental process |
| 8.13E-17 | 5620 | 4583 | 2125 | 13797 | GO:0050896 | GO:BP | response to stimulus |
| 1.04E-16 | 1465 | 4583 | 651 | 13797 | GO:0009888 | GO:BP | tissue development |
| 1.10E-16 | 1418 | 4583 | 633 | 13797 | GO:0022008 | GO:BP | neurogenesis |
| 1.95E-16 | 2209 | 4583 | 927 | 13797 | GO:0048468 | GO:BP | cell development |
| 3.90E-16 | 987 | 4583 | 464 | 13797 | GO:0048666 | GO:BP | neuron development |
| 5.43E-16 | 2117 | 4583 | 891 | 13797 | GO:0051239 | GO:BP | regulation of multicellular organismal process |
| 5.78E-15 | 831 | 4583 | 398 | 13797 | GO:0000902 | GO:BP | cell morphogenesis |
| 8.75E-15 | 1227 | 4583 | 552 | 13797 | GO:0048699 | GO:BP | generation of neurons |
| 1.13E-14 | 821 | 4583 | 393 | 13797 | GO:0035295 | GO:BP | tube development |
| 1.52E-14 | 4049 | 4583 | 1571 | 13797 | GO:0007154 | GO:BP | cell communication |
| 1.61E-14 | 2537 | 4583 | 1036 | 13797 | GO:0042221 | GO:BP | response to chemical |
| 2.59E-14 | 867 | 4583 | 410 | 13797 | GO:0072359 | GO:BP | circulatory system development |
| 4.75E-14 | 572 | 4583 | 290 | 13797 | GO:0048858 | GO:BP | cell projection morphogenesis |
| 5.46E-14 | 3979 | 4583 | 1543 | 13797 | GO:0023052 | GO:BP | signaling |
| 5.81E-14 | 1176 | 4583 | 529 | 13797 | GO:0045595 | GO:BP | regulation of cell differentiation |
| 8.92E-14 | 1165 | 4583 | 524 | 13797 | GO:0030182 | GO:BP | neuron differentiation |
| 9.42E-14 | 1241 | 4583 | 553 | 13797 | GO:1901700 | GO:BP | response to oxygen-containing compound |
| 2.18E-13 | 1903 | 4583 | 799 | 13797 | GO:0010033 | GO:BP | response to organic substance |
| 3.23E-13 | 566 | 4583 | 285 | 13797 | GO:0120039 | GO:BP | plasma membrane bounded cell projection morphogenesis |

| | | | | | | | |
|---|---|---|---|---|---|---|---|
| 4.89E-13 | 1082 | 4583 | 489 | 13797 | GO:0016477 | GO:BP | cell migration |
| 4.98E-13 | 4751 | 4583 | 1803 | 13797 | GO:0051716 | GO:BP | cellular response to stimulus |
| 9.53E-13 | 1242 | 4583 | 549 | 13797 | GO:0009719 | GO:BP | response to endogenous stimulus |
| 1.04E-12 | 698 | 4583 | 337 | 13797 | GO:0007167 | GO:BP | enzyme-linked receptor protein signaling pathway |
| 1.19E-12 | 701 | 4583 | 338 | 13797 | GO:0030334 | GO:BP | regulation of cell migration |
| 1.45E-12 | 657 | 4583 | 320 | 13797 | GO:0035239 | GO:BP | tube morphogenesis |
| 1.72E-12 | 552 | 4583 | 277 | 13797 | GO:0048812 | GO:BP | neuron projection morphogenesis |
| 2.05E-12 | 3645 | 4583 | 1416 | 13797 | GO:0007165 | GO:BP | signal transduction |
| 3.30E-12 | 7557 | 4583 | 2738 | 13797 | GO:0050789 | GO:BP | regulation of biological process |
| 4.95E-12 | 739 | 4583 | 351 | 13797 | GO:2000145 | GO:BP | regulation of cell motility |
| 5.64E-12 | 652 | 4583 | 316 | 13797 | GO:0032989 | GO:BP | cellular anatomical entity morphogenesis |
| 5.68E-12 | 1065 | 4583 | 478 | 13797 | GO:2000026 | GO:BP | regulation of multicellular organismal development |
| 6.11E-12 | 882 | 4583 | 407 | 13797 | GO:0060429 | GO:BP | epithelium development |
| 1.01E-11 | 7800 | 4583 | 2814 | 13797 | GO:0065007 | GO:BP | biological regulation |
| 1.05E-11 | 558 | 4583 | 277 | 13797 | GO:0030155 | GO:BP | regulation of cell adhesion |
| 1.12E-11 | 770 | 4583 | 362 | 13797 | GO:0040012 | GO:BP | regulation of locomotion |
| 1.30E-11 | 2088 | 4583 | 857 | 13797 | GO:0065008 | GO:BP | regulation of biological quality |
| 1.43E-11 | 832 | 4583 | 386 | 13797 | GO:0051241 | GO:BP | negative regulation of multicellular organismal process |
| 3.34E-11 | 7213 | 4583 | 2617 | 13797 | GO:0050794 | GO:BP | regulation of cellular process |
| 3.98E-11 | 2498 | 4583 | 1002 | 13797 | GO:0010646 | GO:BP | regulation of cell communication |
| 4.32E-11 | 2507 | 4583 | 1005 | 13797 | GO:0023051 | GO:BP | regulation of signaling |
| 6.21E-11 | 4475 | 4583 | 1693 | 13797 | GO:0048518 | GO:BP | positive regulation of biological process |
| 7.26E-11 | 1130 | 4583 | 498 | 13797 | GO:0042127 | GO:BP | regulation of cell population proliferation |
| 1.05E-10 | 278 | 4583 | 156 | 13797 | GO:0031589 | GO:BP | cell-substrate adhesion |
| 1.26E-10 | 216 | 4583 | 128 | 13797 | GO:0045229 | GO:BP | external encapsulating structure organization |
| 1.26E-10 | 216 | 4583 | 128 | 13797 | GO:0030198 | GO:BP | extracellular matrix organization |
| 1.33E-10 | 870 | 4583 | 397 | 13797 | GO:1901701 | GO:BP | cellular response to oxygen-containing compound |
| 1.61E-10 | 3942 | 4583 | 1506 | 13797 | GO:0048519 | GO:BP | negative regulation of biological process |
| 2.76E-10 | 1379 | 4583 | 589 | 13797 | GO:0008283 | GO:BP | cell population proliferation |
| 3.40E-10 | 218 | 4583 | 128 | 13797 | GO:0043062 | GO:BP | extracellular structure organization |
| 3.67E-10 | 531 | 4583 | 261 | 13797 | GO:0001568 | GO:BP | blood vessel development |
| 4.03E-10 | 556 | 4583 | 271 | 13797 | GO:0001944 | GO:BP | vasculature development |
| 4.20E-10 | 4136 | 4583 | 1570 | 13797 | GO:0048522 | GO:BP | positive regulation of cellular process |
| 4.54E-10 | 517 | 4583 | 255 | 13797 | GO:0048667 | GO:BP | cell morphogenesis involved in neuron differentiation |
| 4.90E-10 | 1061 | 4583 | 468 | 13797 | GO:0071495 | GO:BP | cellular response to endogenous stimulus |
| 6.98E-10 | 414 | 4583 | 212 | 13797 | GO:0061564 | GO:BP | axon development |
| 7.29E-10 | 1251 | 4583 | 539 | 13797 | GO:0048870 | GO:BP | cell motility |
| 7.75E-10 | 546 | 4583 | 266 | 13797 | GO:0070848 | GO:BP | response to growth factor |
| 8.09E-10 | 732 | 4583 | 340 | 13797 | GO:0009887 | GO:BP | animal organ morphogenesis |

| | | | | | | | |
|---|---|---|---|---|---|---|---|
| 9.43E-10 | 1025 | 4583 | 453 | 13797 | GO:0051094 | GO:BP | positive regulation of developmental process |
| 1.04E-09 | 1269 | 4583 | 545 | 13797 | GO:0141124 | GO:BP | intracellular signaling cassette |
| 1.42E-09 | 881 | 4583 | 397 | 13797 | GO:0048646 | GO:BP | anatomical structure formation involved in morphogenesis |
| 1.65E-09 | 861 | 4583 | 389 | 13797 | GO:0040011 | GO:BP | locomotion |
| 1.80E-09 | 3731 | 4583 | 1425 | 13797 | GO:0048523 | GO:BP | negative regulation of cellular process |
| 2.29E-09 | 1857 | 4583 | 760 | 13797 | GO:0070887 | GO:BP | cellular response to chemical stimulus |
| 2.30E-09 | 189 | 4583 | 113 | 13797 | GO:0097485 | GO:BP | neuron projection guidance |
| 2.30E-09 | 189 | 4583 | 113 | 13797 | GO:0007411 | GO:BP | axon guidance |
| 3.76E-09 | 527 | 4583 | 256 | 13797 | GO:0071363 | GO:BP | cellular response to growth factor stimulus |
| 4.79E-09 | 2161 | 4583 | 868 | 13797 | GO:0009966 | GO:BP | regulation of signal transduction |
| 6.13E-09 | 1395 | 4583 | 588 | 13797 | GO:0071310 | GO:BP | cellular response to organic substance |
| 6.62E-09 | 685 | 4583 | 318 | 13797 | GO:0022603 | GO:BP | regulation of anatomical structure morphogenesis |
| 9.28E-09 | 689 | 4583 | 319 | 13797 | GO:0045597 | GO:BP | positive regulation of cell differentiation |
| 1.15E-08 | 1491 | 4583 | 622 | 13797 | GO:0008219 | GO:BP | cell death |
| 1.15E-08 | 1491 | 4583 | 622 | 13797 | GO:0012501 | GO:BP | programmed cell death |
| 2.03E-08 | 2765 | 4583 | 1079 | 13797 | GO:0048583 | GO:BP | regulation of response to stimulus |
| 2.60E-08 | 455 | 4583 | 224 | 13797 | GO:0048514 | GO:BP | blood vessel morphogenesis |
| 2.60E-08 | 640 | 4583 | 298 | 13797 | GO:0033993 | GO:BP | response to lipid |
| 3.51E-08 | 1325 | 4583 | 558 | 13797 | GO:0120036 | GO:BP | plasma membrane bounded cell projection organization |
| 3.70E-08 | 606 | 4583 | 284 | 13797 | GO:0034330 | GO:BP | cell junction organization |
| 4.71E-08 | 776 | 4583 | 350 | 13797 | GO:0040007 | GO:BP | growth |
| 5.63E-08 | 1355 | 4583 | 568 | 13797 | GO:0030030 | GO:BP | cell projection organization |
| 6.68E-08 | 1259 | 4583 | 532 | 13797 | GO:0048585 | GO:BP | negative regulation of response to stimulus |
| 7.41E-08 | 1099 | 4583 | 472 | 13797 | GO:0010648 | GO:BP | negative regulation of cell communication |
| 9.87E-08 | 1098 | 4583 | 471 | 13797 | GO:0023057 | GO:BP | negative regulation of signaling |
| 1.24E-07 | 698 | 4583 | 318 | 13797 | GO:0051093 | GO:BP | negative regulation of developmental process |
| 1.76E-07 | 255 | 4583 | 138 | 13797 | GO:0072001 | GO:BP | renal system development |
| 2.22E-07 | 372 | 4583 | 187 | 13797 | GO:0007409 | GO:BP | axonogenesis |
| 2.72E-07 | 1179 | 4583 | 499 | 13797 | GO:0043067 | GO:BP | regulation of programmed cell death |
| 2.80E-07 | 481 | 4583 | 231 | 13797 | GO:0008285 | GO:BP | negative regulation of cell population proliferation |
| 3.43E-07 | 677 | 4583 | 308 | 13797 | GO:0098657 | GO:BP | import into cell |
| 3.63E-07 | 285 | 4583 | 150 | 13797 | GO:0001503 | GO:BP | ossification |
| 3.67E-07 | 1438 | 4583 | 594 | 13797 | GO:0006915 | GO:BP | apoptotic process |
| 4.24E-07 | 1171 | 4583 | 495 | 13797 | GO:0051240 | GO:BP | positive regulation of multicellular organismal process |
| 4.76E-07 | 246 | 4583 | 133 | 13797 | GO:0001822 | GO:BP | kidney development |
| 5.36E-07 | 419 | 4583 | 205 | 13797 | GO:2000147 | GO:BP | positive regulation of cell motility |
| 5.83E-07 | 407 | 4583 | 200 | 13797 | GO:0030335 | GO:BP | positive regulation of cell migration |
| 8.72E-07 | 1189 | 4583 | 500 | 13797 | GO:0003008 | GO:BP | system process |
| 9.60E-07 | 431 | 4583 | 209 | 13797 | GO:0040017 | GO:BP | positive regulation of locomotion |

| | | | | | | | |
|---|---|---|---|---|---|---|---|
| 1.11E-06 | 434 | 4583 | 210 | 13797 | GO:0007169 | GO:BP | transmembrane receptor protein tyrosine kinase signaling pathway |
| 1.59E-06 | 661 | 4583 | 299 | 13797 | GO:0060284 | GO:BP | regulation of cell development |
| 1.97E-06 | 1100 | 4583 | 465 | 13797 | GO:0007267 | GO:BP | cell-cell signaling |
| 1.97E-06 | 1009 | 4583 | 431 | 13797 | GO:0009968 | GO:BP | negative regulation of signal transduction |
| 2.36E-06 | 417 | 4583 | 202 | 13797 | GO:0051960 | GO:BP | regulation of nervous system development |
| 2.48E-06 | 586 | 4583 | 269 | 13797 | GO:0048589 | GO:BP | developmental growth |
| 2.53E-06 | 1702 | 4583 | 685 | 13797 | GO:0009605 | GO:BP | response to external stimulus |
| 2.72E-06 | 661 | 4583 | 298 | 13797 | GO:0014070 | GO:BP | response to organic cyclic compound |
| 3.12E-06 | 381 | 4583 | 187 | 13797 | GO:0001525 | GO:BP | angiogenesis |
| 4.28E-06 | 812 | 4583 | 355 | 13797 | GO:1901698 | GO:BP | response to nitrogen compound |
| 5.29E-06 | 1144 | 4583 | 479 | 13797 | GO:0042981 | GO:BP | regulation of apoptotic process |
| 6.48E-06 | 575 | 4583 | 263 | 13797 | GO:0098609 | GO:BP | cell-cell adhesion |
| 6.86E-06 | 347 | 4583 | 172 | 13797 | GO:0045785 | GO:BP | positive regulation of cell adhesion |
| 7.08E-06 | 352 | 4583 | 174 | 13797 | GO:0050767 | GO:BP | regulation of neurogenesis |
| 9.37E-06 | 485 | 4583 | 227 | 13797 | GO:0030855 | GO:BP | epithelial cell differentiation |
| 9.46E-06 | 1924 | 4583 | 761 | 13797 | GO:0035556 | GO:BP | intracellular signal transduction |
| 1.07E-05 | 149 | 4583 | 87 | 13797 | GO:0001763 | GO:BP | morphogenesis of a branching structure |
| 1.32E-05 | 522 | 4583 | 241 | 13797 | GO:0000165 | GO:BP | MAPK cascade |
| 1.62E-05 | 500 | 4583 | 232 | 13797 | GO:0045596 | GO:BP | negative regulation of cell differentiation |
| 1.74E-05 | 1988 | 4583 | 782 | 13797 | GO:0051128 | GO:BP | regulation of cellular component organization |
| 1.90E-05 | 410 | 4583 | 196 | 13797 | GO:0050808 | GO:BP | synapse organization |
| 2.38E-05 | 381 | 4583 | 184 | 13797 | GO:0010817 | GO:BP | regulation of hormone levels |
| 2.97E-05 | 1278 | 4583 | 524 | 13797 | GO:0023056 | GO:BP | positive regulation of signaling |
| 4.54E-05 | 376 | 4583 | 181 | 13797 | GO:0001501 | GO:BP | skeletal system development |
| 5.20E-05 | 752 | 4583 | 327 | 13797 | GO:0010243 | GO:BP | response to organonitrogen compound |
| 5.28E-05 | 1274 | 4583 | 521 | 13797 | GO:0010647 | GO:BP | positive regulation of cell communication |
| 6.54E-05 | 171 | 4583 | 95 | 13797 | GO:0010810 | GO:BP | regulation of cell-substrate adhesion |
| 9.06E-05 | 138 | 4583 | 80 | 13797 | GO:0061138 | GO:BP | morphogenesis of a branching epithelium |
| 0.000112591 | 450 | 4583 | 209 | 13797 | GO:0010975 | GO:BP | regulation of neuron projection development |
| 0.000157704 | 341 | 4583 | 165 | 13797 | GO:0048732 | GO:BP | gland development |
| 0.000176435 | 1220 | 4583 | 498 | 13797 | GO:0042592 | GO:BP | homeostatic process |
| 0.000178498 | 693 | 4583 | 302 | 13797 | GO:0032101 | GO:BP | regulation of response to external stimulus |
| 0.000189242 | 1554 | 4583 | 619 | 13797 | GO:0048584 | GO:BP | positive regulation of response to stimulus |
| 0.000195887 | 176 | 4583 | 96 | 13797 | GO:0007160 | GO:BP | cell-matrix adhesion |
| 0.000272956 | 368 | 4583 | 175 | 13797 | GO:0009611 | GO:BP | response to wounding |
| 0.000286822 | 898 | 4583 | 378 | 13797 | GO:0006468 | GO:BP | protein phosphorylation |
| 0.000302734 | 391 | 4583 | 184 | 13797 | GO:1901652 | GO:BP | response to peptide |
| 0.00033722 | 159 | 4583 | 88 | 13797 | GO:0060348 | GO:BP | bone development |
| 0.000356131 | 527 | 4583 | 237 | 13797 | GO:0006897 | GO:BP | endocytosis |
| 0.000470475 | 223 | 4583 | 115 | 13797 | GO:0060485 | GO:BP | mesenchyme development |

| | | | | | | | |
|---|---|---|---|---|---|---|---|
| 0.000629984 | 450 | 4583 | 206 | 13797 | GO:0043408 | GO:BP | regulation of MAPK cascade |
| 0.000639785 | 468 | 4583 | 213 | 13797 | GO:0048729 | GO:BP | tissue morphogenesis |
| 0.000660536 | 354 | 4583 | 168 | 13797 | GO:0007264 | GO:BP | small GTPase-mediated signal transduction |
| 0.000697927 | 42 | 4583 | 32 | 13797 | GO:0085029 | GO:BP | extracellular matrix assembly |
| 0.000736313 | 709 | 4583 | 305 | 13797 | GO:0001775 | GO:BP | cell activation |
| 0.000742035 | 717 | 4583 | 308 | 13797 | GO:0043069 | GO:BP | negative regulation of programmed cell death |
| 0.000802958 | 1111 | 4583 | 454 | 13797 | GO:0009967 | GO:BP | positive regulation of signal transduction |
| 0.000807795 | 500 | 4583 | 225 | 13797 | GO:1901699 | GO:BP | cellular response to nitrogen compound |
| 0.000807795 | 500 | 4583 | 225 | 13797 | GO:0040008 | GO:BP | regulation of growth |
| 0.000829315 | 641 | 4583 | 279 | 13797 | GO:0009725 | GO:BP | response to hormone |
| 0.000831079 | 103 | 4583 | 62 | 13797 | GO:0072073 | GO:BP | kidney epithelium development |
| 0.000849913 | 350 | 4583 | 166 | 13797 | GO:0003013 | GO:BP | circulatory system process |
| 0.000921778 | 1082 | 4583 | 443 | 13797 | GO:0016310 | GO:BP | phosphorylation |
| 0.000999425 | 447 | 4583 | 204 | 13797 | GO:0006954 | GO:BP | inflammatory response |
| 0.001141632 | 183 | 4583 | 97 | 13797 | GO:0048762 | GO:BP | mesenchymal cell differentiation |
| 0.001149623 | 146 | 4583 | 81 | 13797 | GO:0001649 | GO:BP | osteoblast differentiation |
| 0.001243586 | 226 | 4583 | 115 | 13797 | GO:0006790 | GO:BP | sulfur compound metabolic process |
| 0.001271498 | 188 | 4583 | 99 | 13797 | GO:0051896 | GO:BP | regulation of phosphatidylinositol 3-kinase/protein kinase B signal transduction |
| 0.001393979 | 104 | 4583 | 62 | 13797 | GO:0044272 | GO:BP | sulfur compound biosynthetic process |
| 0.001632763 | 184 | 4583 | 97 | 13797 | GO:0060541 | GO:BP | respiratory system development |
| 0.001901601 | 2416 | 4583 | 917 | 13797 | GO:0031325 | GO:BP | positive regulation of cellular metabolic process |
| 0.002125196 | 166 | 4583 | 89 | 13797 | GO:0030323 | GO:BP | respiratory tube development |
| 0.002137753 | 77 | 4583 | 49 | 13797 | GO:0032963 | GO:BP | collagen metabolic process |
| 0.002313502 | 387 | 4583 | 179 | 13797 | GO:0002009 | GO:BP | morphogenesis of an epithelium |
| 0.00234171 | 1218 | 4583 | 490 | 13797 | GO:0051049 | GO:BP | regulation of transport |
| 0.002410661 | 869 | 4583 | 362 | 13797 | GO:0009628 | GO:BP | response to abiotic stimulus |
| 0.002468389 | 490 | 4583 | 219 | 13797 | GO:0007507 | GO:BP | heart development |
| 0.002784831 | 101 | 4583 | 60 | 13797 | GO:0045667 | GO:BP | regulation of osteoblast differentiation |
| 0.00293527 | 728 | 4583 | 309 | 13797 | GO:0007417 | GO:BP | central nervous system development |
| 0.00300645 | 200 | 4583 | 103 | 13797 | GO:0007162 | GO:BP | negative regulation of cell adhesion |
| 0.00302635 | 263 | 4583 | 129 | 13797 | GO:0060562 | GO:BP | epithelial tube morphogenesis |
| 0.003057037 | 424 | 4583 | 193 | 13797 | GO:0002521 | GO:BP | leukocyte differentiation |
| 0.003448504 | 628 | 4583 | 271 | 13797 | GO:0008284 | GO:BP | positive regulation of cell population proliferation |
| 0.003545443 | 417 | 4583 | 190 | 13797 | GO:0016049 | GO:BP | cell growth |
| 0.003666871 | 505 | 4583 | 224 | 13797 | GO:0007610 | GO:BP | behavior |
| 0.003841622 | 326 | 4583 | 154 | 13797 | GO:0008015 | GO:BP | blood circulation |
| 0.003940394 | 163 | 4583 | 87 | 13797 | GO:0030324 | GO:BP | lung development |
| 0.003985583 | 115 | 4583 | 66 | 13797 | GO:0048754 | GO:BP | branching morphogenesis of an epithelial tube |
| 0.004076475 | 225 | 4583 | 113 | 13797 | GO:0043491 | GO:BP | phosphatidylinositol 3-kinase/protein kinase B signal transduction |

| | | | | | | | |
|---|---|---|---|---|---|---|---|
| 0.004087823 | 4055 | 4583 | 1480 | 13797 | GO:0031323 | GO:BP | regulation of cellular metabolic process |
| 0.004626968 | 2658 | 4583 | 998 | 13797 | GO:0009893 | GO:BP | positive regulation of metabolic process |
| 0.004637332 | 267 | 4583 | 130 | 13797 | GO:0051962 | GO:BP | positive regulation of nervous system development |
| 0.004869663 | 332 | 4583 | 156 | 13797 | GO:0001667 | GO:BP | ameboidal-type cell migration |
| 0.005095409 | 411 | 4583 | 187 | 13797 | GO:0071396 | GO:BP | cellular response to lipid |
| 0.005466247 | 109 | 4583 | 63 | 13797 | GO:0072006 | GO:BP | nephron development |
| 0.006013982 | 83 | 4583 | 51 | 13797 | GO:0045446 | GO:BP | endothelial cell differentiation |
| 0.006327556 | 555 | 4583 | 242 | 13797 | GO:0001932 | GO:BP | regulation of protein phosphorylation |
| 0.006514184 | 456 | 4583 | 204 | 13797 | GO:0071417 | GO:BP | cellular response to organonitrogen compound |
| 0.00657881 | 693 | 4583 | 294 | 13797 | GO:0043066 | GO:BP | negative regulation of apoptotic process |
| 0.007439478 | 103 | 4583 | 60 | 13797 | GO:0031214 | GO:BP | biomineral tissue development |
| 0.008023446 | 90 | 4583 | 54 | 13797 | GO:0051384 | GO:BP | response to glucocorticoid |
| 0.00877105 | 257 | 4583 | 125 | 13797 | GO:0016358 | GO:BP | dendrite development |
| 0.008873713 | 135 | 4583 | 74 | 13797 | GO:0051897 | GO:BP | positive regulation of phosphatidylinositol 3-kinase/protein kinase B signal transduction |
| 0.009176219 | 142 | 4583 | 77 | 13797 | GO:0042445 | GO:BP | hormone metabolic process |
| 0.009280097 | 282 | 4583 | 135 | 13797 | GO:0042063 | GO:BP | gliogenesis |
| 0.009339429 | 1814 | 4583 | 699 | 13797 | GO:0006796 | GO:BP | phosphate-containing compound metabolic process |
| 0.009463315 | 350 | 4583 | 162 | 13797 | GO:0048568 | GO:BP | embryonic organ development |
| 0.009582299 | 963 | 4583 | 393 | 13797 | GO:0006952 | GO:BP | defense response |
| 0.010189098 | 168 | 4583 | 88 | 13797 | GO:0002573 | GO:BP | myeloid leukocyte differentiation |
| 0.010199394 | 592 | 4583 | 255 | 13797 | GO:0048871 | GO:BP | multicellular organismal-level homeostasis |
| 0.010256112 | 43 | 4583 | 31 | 13797 | GO:1903053 | GO:BP | regulation of extracellular matrix organization |
| 0.010433667 | 161 | 4583 | 85 | 13797 | GO:1901654 | GO:BP | response to ketone |
| 0.010504912 | 65 | 4583 | 42 | 13797 | GO:0060993 | GO:BP | kidney morphogenesis |
| 0.013230268 | 102 | 4583 | 59 | 13797 | GO:0031960 | GO:BP | response to corticosteroid |
| 0.015450701 | 284 | 4583 | 135 | 13797 | GO:0007178 | GO:BP | transmembrane receptor protein serine/threonine kinase signaling pathway |
| 0.015493245 | 388 | 4583 | 176 | 13797 | GO:0010720 | GO:BP | positive regulation of cell development |
| 0.01599667 | 215 | 4583 | 107 | 13797 | GO:0071559 | GO:BP | response to transforming growth factor beta |
| 0.017213079 | 474 | 4583 | 209 | 13797 | GO:0048598 | GO:BP | embryonic morphogenesis |
| 0.017821405 | 1832 | 4583 | 703 | 13797 | GO:0006793 | GO:BP | phosphorus metabolic process |
| 0.018181612 | 213 | 4583 | 106 | 13797 | GO:0010721 | GO:BP | negative regulation of cell development |
| 0.021366908 | 405 | 4583 | 182 | 13797 | GO:0010035 | GO:BP | response to inorganic substance |
| 0.02141962 | 950 | 4583 | 386 | 13797 | GO:0051130 | GO:BP | positive regulation of cellular component organization |
| 0.022550337 | 626 | 4583 | 266 | 13797 | GO:0030029 | GO:BP | actin filament-based process |
| 0.02290501 | 168 | 4583 | 87 | 13797 | GO:0031099 | GO:BP | regeneration |
| 0.025917772 | 675 | 4583 | 284 | 13797 | GO:0140352 | GO:BP | export from cell |
| 0.030161416 | 317 | 4583 | 147 | 13797 | GO:0050673 | GO:BP | epithelial cell proliferation |
| 0.031470872 | 210 | 4583 | 104 | 13797 | GO:0071560 | GO:BP | cellular response to transforming growth factor beta stimulus |

| | | | | | | | |
|---|---|---|---|---|---|---|---|
| 0.031597838 | 335 | 4583 | 154 | 13797 | GO:0043434 | GO:BP | response to peptide hormone |
| 0.032770754 | 277 | 4583 | 131 | 13797 | GO:1901653 | GO:BP | cellular response to peptide |
| 0.03455483 | 610 | 4583 | 259 | 13797 | GO:0031344 | GO:BP | regulation of cell projection organization |
| 0.03506258 | 1266 | 4583 | 499 | 13797 | GO:0044281 | GO:BP | small molecule metabolic process |
| 0.035666961 | 10947 | 4583 | 3744 | 13797 | GO:0009987 | GO:BP | cellular process |
| 0.035751011 | 67 | 4583 | 42 | 13797 | GO:0001823 | GO:BP | mesonephros development |
| 0.03605675 | 295 | 4583 | 138 | 13797 | GO:0009410 | GO:BP | response to xenobiotic stimulus |
| 0.038090618 | 65 | 4583 | 41 | 13797 | GO:0072164 | GO:BP | mesonephric tubule development |
| 0.038090618 | 65 | 4583 | 41 | 13797 | GO:0072163 | GO:BP | mesonephric epithelium development |
| 0.038423229 | 531 | 4583 | 229 | 13797 | GO:0061061 | GO:BP | muscle structure development |
| 0.039228797 | 1544 | 4583 | 598 | 13797 | GO:0032879 | GO:BP | regulation of localization |
| 0.040415019 | 63 | 4583 | 40 | 13797 | GO:0001656 | GO:BP | metanephros development |
| 0.040424542 | 233 | 4583 | 113 | 13797 | GO:0050769 | GO:BP | positive regulation of neurogenesis |
| 0.044968603 | 1246 | 4583 | 491 | 13797 | GO:1902531 | GO:BP | regulation of intracellular signal transduction |
| 0.047077126 | 241 | 4583 | 116 | 13797 | GO:0001666 | GO:BP | response to hypoxia |
| 3.76E-33 | 3486 | 4583 | 1470 | 13797 | GO:0071944 | GO:CC | cell periphery |
| 1.62E-24 | 3158 | 4583 | 1312 | 13797 | GO:0005886 | GO:CC | plasma membrane |
| 3.03E-14 | 573 | 4583 | 291 | 13797 | GO:0009986 | GO:CC | cell surface |
| 7.75E-14 | 318 | 4583 | 181 | 13797 | GO:0031012 | GO:CC | extracellular matrix |
| 1.89E-13 | 320 | 4583 | 181 | 13797 | GO:0030312 | GO:CC | external encapsulating structure |
| 1.39E-12 | 253 | 4583 | 149 | 13797 | GO:0062023 | GO:CC | collagen-containing extracellular matrix |
| 6.06E-12 | 5861 | 4583 | 2172 | 13797 | GO:0016020 | GO:CC | membrane |
| 7.00E-09 | 975 | 4583 | 430 | 13797 | GO:0098590 | GO:CC | plasma membrane region |
| 1.31E-08 | 1121 | 4583 | 484 | 13797 | GO:0005576 | GO:CC | extracellular region |
| 3.49E-08 | 1695 | 4583 | 694 | 13797 | GO:0030054 | GO:CC | cell junction |
| 1.81E-06 | 729 | 4583 | 325 | 13797 | GO:0005615 | GO:CC | extracellular space |
| 8.33E-06 | 548 | 4583 | 252 | 13797 | GO:0070161 | GO:CC | anchoring junction |
| 8.44E-06 | 83 | 4583 | 56 | 13797 | GO:0005604 | GO:CC | basement membrane |
| 1.81E-05 | 820 | 4583 | 355 | 13797 | GO:0036477 | GO:CC | somatodendritic compartment |
| 0.000629923 | 535 | 4583 | 239 | 13797 | GO:0043025 | GO:CC | neuronal cell body |
| 0.000656582 | 236 | 4583 | 120 | 13797 | GO:0043235 | GO:CC | receptor complex |
| 0.000742241 | 222 | 4583 | 114 | 13797 | GO:0009925 | GO:CC | basal plasma membrane |
| 0.000928037 | 206 | 4583 | 107 | 13797 | GO:0009897 | GO:CC | external side of plasma membrane |
| 0.00104623 | 595 | 4583 | 261 | 13797 | GO:0097447 | GO:CC | dendritic tree |
| 0.001108927 | 11967 | 4583 | 4077 | 13797 | GO:0110165 | GO:CC | cellular anatomical entity |
| 0.001319338 | 231 | 4583 | 117 | 13797 | GO:0045178 | GO:CC | basal part of cell |
| 0.001410459 | 594 | 4583 | 260 | 13797 | GO:0030425 | GO:CC | dendrite |
| 0.001538564 | 198 | 4583 | 103 | 13797 | GO:0016323 | GO:CC | basolateral plasma membrane |
| 0.002651781 | 1142 | 4583 | 462 | 13797 | GO:0043005 | GO:CC | neuron projection |
| 0.002782682 | 421 | 4583 | 192 | 13797 | GO:0098552 | GO:CC | side of membrane |

| | | | | | | | |
|---|---|---|---|---|---|---|---|
| 0.003101305 | 601 | 4583 | 261 | 13797 | GO:0044297 | GO:CC | cell body |
| 0.015174714 | 347 | 4583 | 160 | 13797 | GO:0097060 | GO:CC | synaptic membrane |
| 0.017660374 | 1246 | 4583 | 494 | 13797 | GO:0045202 | GO:CC | synapse |
| 0.032696725 | 599 | 4583 | 255 | 13797 | GO:0030424 | GO:CC | axon |
| 0.034293984 | 397 | 4583 | 178 | 13797 | GO:0005911 | GO:CC | cell-cell junction |
| 5.22E-10 | 7439 | 4583 | 2682 | 13797 | GO:0005515 | GO:MF | protein binding |
| 1.80E-08 | 1715 | 4583 | 703 | 13797 | GO:0042802 | GO:MF | identical protein binding |
| 2.47E-05 | 9993 | 4583 | 3470 | 13797 | GO:0005488 | GO:MF | binding |
| 0.000408964 | 47 | 4583 | 35 | 13797 | GO:0050840 | GO:MF | extracellular matrix binding |
| 0.001008971 | 553 | 4583 | 245 | 13797 | GO:0042803 | GO:MF | protein homodimerization activity |
| 0.001026901 | 48 | 4583 | 35 | 13797 | GO:0005518 | GO:MF | collagen binding |
| 0.001821082 | 98 | 4583 | 59 | 13797 | GO:0005201 | GO:MF | extracellular matrix structural constituent |
| 0.00213736 | 906 | 4583 | 376 | 13797 | GO:0005102 | GO:MF | signaling receptor binding |
| 0.004363245 | 4461 | 4583 | 1618 | 13797 | GO:0036094 | GO:MF | small molecule binding |
| 0.004431625 | 113 | 4583 | 65 | 13797 | GO:0019838 | GO:MF | growth factor binding |
| 0.005929725 | 770 | 4583 | 323 | 13797 | GO:0046983 | GO:MF | protein dimerization activity |
| 0.019857768 | 4330 | 4583 | 1566 | 13797 | GO:0043167 | GO:MF | ion binding |
| 0.026696125 | 207 | 4583 | 103 | 13797 | GO:0050839 | GO:MF | cell adhesion molecule binding |
| 0.044855015 | 59 | 4583 | 38 | 13797 | GO:0019199 | GO:MF | transmembrane receptor protein kinase activity |
